# Leveraging genomic redundancy to improve inference and alignment of orthologous proteins

**DOI:** 10.1101/2023.01.24.525427

**Authors:** Marc Singleton, Michael Eisen

## Abstract

Identifying protein sequences with common ancestry is a core task in bioinformatics and evolutionary biology. However, methods for inferring and aligning such sequences in annotated genomes have not kept pace with the increasing scale and complexity of the available data. Thus, in this work we implemented several improvements to the traditional methodology that more fully leverage the redundancy of closely related genomes and the organization of their annotations. Two highlights include the application of the more flexible *k*-clique percolation algorithm for identifying clusters of orthologous proteins and the development of a novel technique for removing poorly supported regions of alignments with a phylogenetic HMM. In making the latter, we also wrote a fully documented Python package Homomorph that implements standard HMM algorithms and created a set of tutorials to promote its use by a wide audience. We applied the resulting pipeline to a set of 33 annotated *Drosophila* genomes, generating 22,813 orthologous groups and 8,566 high-quality alignments.

## Introduction

Comparative genomics is a powerful tool for yielding insights into evolutionary relationships, molecular function, and the forces that drive gene, genome, and population evolution. These methods often rely on the identification of homologous sequences or homologs, that is sequences with common ancestry, since this ensures that differences between sequences reflect variations in evolution from a common point of divergence. However, many analyses impose the additional condition that the sequences have diverged through speciation events (orthology) rather than duplications (paralogy) or other mechanisms such as horizontal gene transfer [20]. The underlying assumption is “orthologs” have conserved equivalent functions whereas “paralogs,” by virtue of their redundancy, are more likely to diverge [51, 49, 3, 52, 57].^1^ This relationship between orthology and function is an essential component of modern biological research since it permits the transfer of annotations between biological systems using sequence similarity alone.

Given this importance, methods for inferring orthologous groups of proteins were developed shortly after the first genomes were sequenced in the late 1990s [21, 24, 55]. One early and influential approach was to cluster triangles of hits resulting from homology searches between pairs of genomes [58]. Graph-based approaches have remained popular, and in the intervening years many other researchers have refined this method by implementing various pre- and post-processing steps.

Despite these improvements, many databases and pipelines use the same triangle clustering algorithm or other methods which require relatively few hits between sequences to infer an orthologous group, *e.g*. connected components or Markov clustering [53, 17, 38, 30, 40, 16, 61, 10]. However, the scale of biological sequence data has changed dramatically. For example, in the last decade, the number of annotated genomes available from NCBI has increased nearly 20-fold and currently exceeds 900 (Fig. S1). Though this figure is only a rough proxy of the total number of assemblies available, it will likely continue to grow rapidly in the coming years as many large-scale genome assembly efforts such as i5K, the Bird 10,000 Genomes Project, and the Vertebrate Genomes Project have already yielded results [60, 19, 54]. Thus, the dense taxonomic sampling made possible by these projects poses new challenges and opportunities for the standard methods of orthology inference and alignment, which implicitly assume fewer and more distantly related genomes or fail to fully leverage the redundancy and organization of their annotations.

In this work we therefore developed a computational pipeline that can robustly infer and align orthologous groups of proteins even when the genomes are highly redundant. Like many other orthology inference pipelines, our overall approach is based on clustering a graph of hits from homology searches. However, we modified many details to maximize the detection of highly diverged orthologs while also minimizing the impact of incomplete or incorrect annotations. Furthermore, since modern genome annotation pipelines frequently produce gene models and protein sequences in tandem, we implemented an additional clustering step to organize the resulting orthologous groups of proteins into gene-level units. However, most of our efforts were focused on the final step of aligning the orthologous sequences. Though genome annotation pipelines are often proficient at identifying the overall locus of genes, the accurate identification of exon boundaries and start codons when transcript evidence is limited remains an ongoing challenge [22, 12]. Consequently, protein sequences derived from annotation pipelines can include non-homologous segments of significant length or exclude highly conserved segments. Such heterogeneity in the structure and length of the sequences in an orthologous group poses many challenges for their alignment and subsequent analysis. Thus, we implemented several novel quality control and data cleaning steps to correct mis-alignments and identify likely sequencing, assembly, or annotation errors.

To develop these methods, we chose a set of 33 assembled and annotated *Drosophila* genomes, which includes all 12 species from the original *Drosophila* 12 Genomes Consortium [9]. However, the genomes of these 12 species have been re-sequenced since their first release, which has resulted in substantial improvements in their assemblies and annotations. Despite these developments and other several other recent genome assembly projects of species in the *Drosophila* genus, there is not yet a collection of high-quality alignments of orthologous proteins that reflects these improvements in genome assembly and diversity [43, 34]. Given the *Drosophila* genus spans diverse habitats and over 50 million years of evolution but maintains a conserved life cycle and body plan, such a resource would facilitate a new generation of studies that illuminate the forces that drive protein evolution in unprecedented detail [63, 50].

## Results

### Pipeline overview

Our pipeline follows a similar overall approach to other graph-based methods of orthology inference. First, protein sequences from annotated genomes are collected (Fig. 1A), and homology searches are then conducted between all query-target pairs of genomes (Fig. 1B). The raw output from these homology searches is processed to yield best hits between pairs of sequences (Fig. 1C). Next, the network of best hits is clustered into self-consistent orthologous groups (Fig. 1D). Since genes can have multiplied associated isoforms, we then implemented a novel second clustering step where orthologous groups are grouped by their parent genes, which are represented by the two sets of clusters with warm and cool colors, respectively (Fig. 1E). Finally, representative sequences in each orthologous group are aligned (Fig. 1F). In the following sections, we discuss each of these and other steps which were omitted for clarity in greater detail.

**Fig 1.**
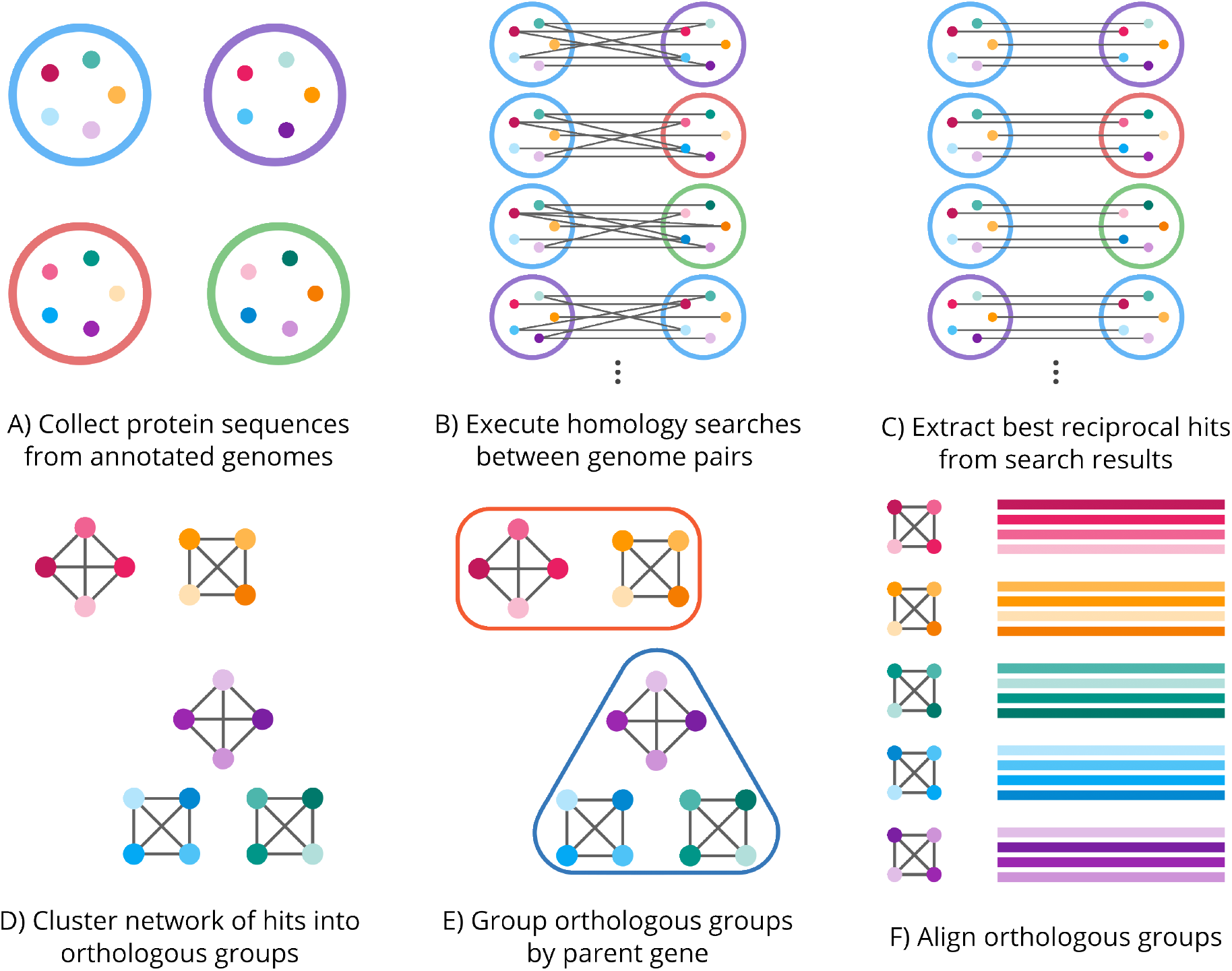
Overview of homology inference pipeline. **(A-C)** The large, open circles represent the annotated genomes, and the small, filled circles represent the protein sequences associated with each annotation. Sequences that share homology are colored with shades of the same hue. **(D-F)** The small, filled circles and lines represent the same sequences and best hits from the previous steps.

### Input genomes and pre-processing

All annotated genomes in the genus *Drosophila* available in April 2022 were downloaded from NCBI’s RefSeq database. The assemblies annotated by the NCBI eukaryotic genome annotation pipeline have passed several quality checks and all have supporting transcript evidence, so the annotations are generally highly complete [59]. The *D. miranda* annotation was excluded due to its unusual karyotype [11]. Other annotations were excluded after preliminary clustering showed a deficiency in the number of orthologous groups containing those genomes, indicating their annotations were less complete (data not shown). The *D. melanogaster* annotation was downloaded from FlyBase [25]. In total, the input data consists of 33 genomes, which are listed in Table S1. Many genes have transcripts that differ only in their UTRs, and as a result there are many duplicate protein sequences in the annotations. Though not strictly necessary, we removed the duplicates in our pipeline, which greatly reduced the computational burden of later steps.

### Extraction of best hits from BLAST output

The protein sequences in each genome annotation were searched against each other in reciprocal pairs using BLAST, yielding a list of high-scoring segment pairs (HSPs) for each query-target pair [5]. HSPs are local alignments, meaning they do not necessarily span the entire lengths of the query and target sequences. Consequently, the search algorithm may return multiple HSPs for each query-target pair if statistically significant regions of homology are separated by nonhomologous or poorly conserved regions. Though the most significant HSP is often used to represent all HSPs between a query-target pair, this approach can fail to rank the pairs by their overall significance if their alignments are broken into multiple HSPs. Furthermore, since query-target pairs were later filtered by the amount overlap between their sequences, it can also exclude pairs that pass the overlap threshold even if the most significant HSP alone does not. Thus, HSPs were merged into a single object called a hit. The best hits for each query were then taken as the highest-scoring hits that passed a minimum overlap criterion and were reciprocal between the query and target sequences.

### Clustering in orthologous groups

The best hits between sequences are naturally visualized as a graph where sequences are nodes and best hits are edges between nodes. Two connected components, sets of nodes joined by a sequence of edges, are shown (Fig. 2A,B). The sequences in the first (Fig. 2A) all contain C2H2 zinc fingers, whereas the sequences in the second (Fig. 2B) are members of the Par-1 family of serine/threonine protein kinases. In both components, some sets of nodes have a high density of edges, forming distinct clusters, whereas other nodes are only sparsely connected to their neighbors. To better understand the structure of these two components, we calculated the number of sequences, unique genes, and unique species in each. The first has 385, 346, and 33 sequences, genes, and species, respectively, and the second has 222, 33, and 33 sequences, genes, and species, respectively. We then plotted the relationship between the number sequences and unique genes across all components to see if this pattern holds true generally (Fig. 2C). Two distinct trendlines are apparent. The first increases linearly with the number of sequences with a slope of one, indicating each sequence is generally associated with a unique gene. The second is constant with an intercept of 33, indicating the number of unique genes quickly saturates at the total number of genomes. Thus, there are generally two classes of components. The first is composed of many distinct genes, whereas the second is composed of many different isoforms of a single group of genes.

**Fig 2.**
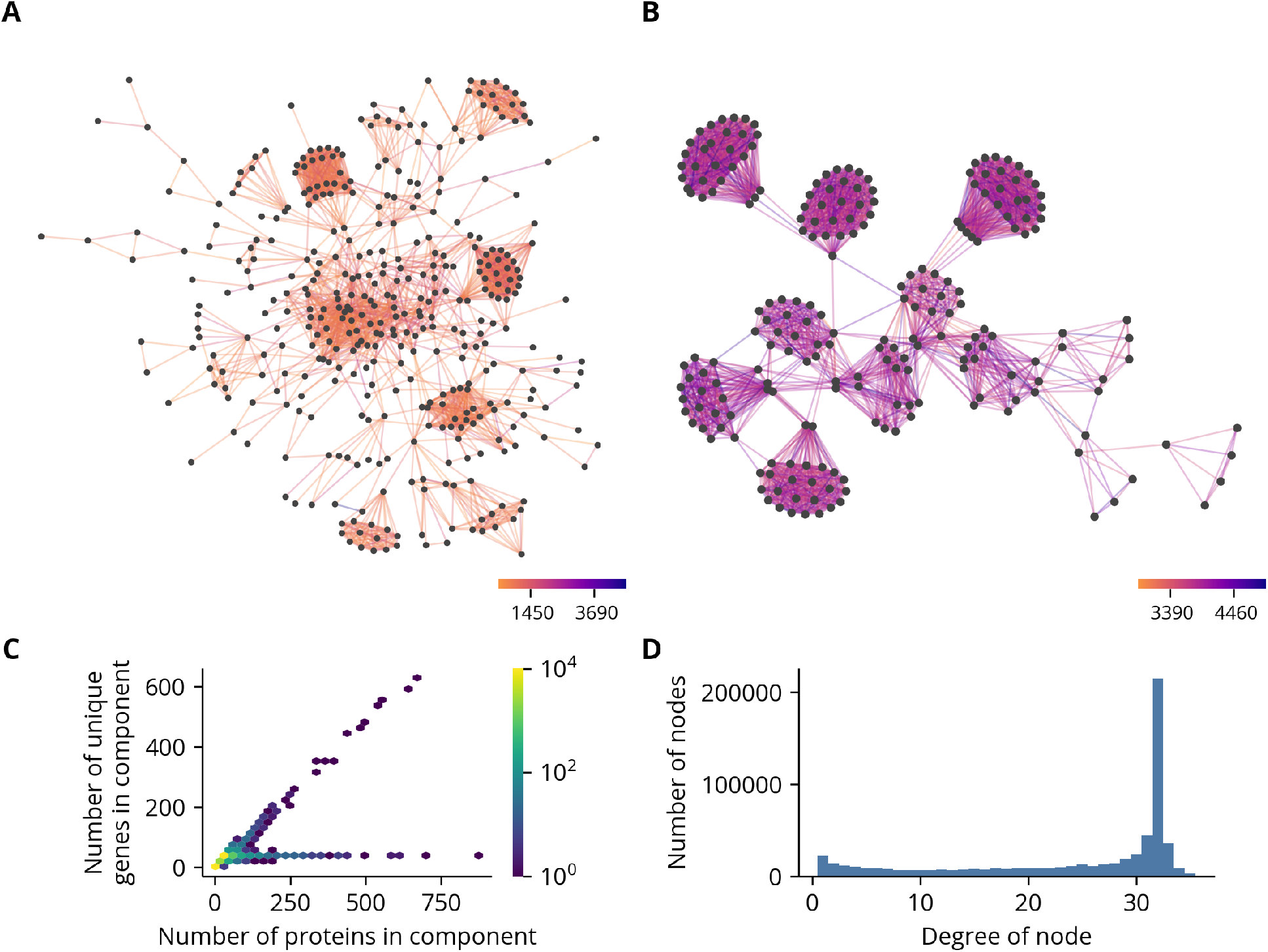
Selected connected components of hit graph and summary statistics. **(A-B)** Two distinct connected components of the hit graph. Edges are colored by the value of their bit score. **(C)** Hexbin plot of the number of sequences and the number of unique genes in each component. **(D)** Histogram of number edges associated with each sequence, *i.e*. the degree of each node. Only the lower 99th percentile of the distribution is shown.

The diffuse networks observed in the first component class are likely the result of a combination of factors, including rapid evolution, gene duplication, and annotation errors. Regardless of their origin, these hits are not strong candidates for comparative analyses since an orthology relationship is supported by relatively few genome pairs. Instead, likely orthologs should consistently identify each other as best reciprocal hits across many genome pairs. The same is true of the hits in the second component class. Although the genes as a unit form a single orthologous group, sequences with few hits are likely non-conserved or tissue-specific isoforms. Thus, orthologous groups can be operationally defined as self-consistent clusters in the hit graph. However, sequence divergence or assembly and annotation errors may prevent a best reciprocal hit between orthologous sequences across all genome pairs. In fact, although the most common number of reciprocal hits is 32, one fewer than the total number of genomes, many sequences have fewer (Fig. 2D). Thus, the clustering method should require a high degree of self-consistency without demanding complete consensus.

The identification of sets of densely connected nodes in graphs is known as community detection in network analysis. While many community detection algorithms are available, only some are commonly used in the context of orthology inference. One early method that remains popular is building clusters progressively by identifying nodes that form a triangle with at least two other nodes in the cluster [58, 30]. Other approaches include the MCL algorithm, which clusters graphs by similating stochastic flow, and connected components or other single-linkage criteria [53, 17, 38, 16, 61, 10]. While these methods are robust when clustering hit graphs derived from smaller or more diverse sets of genomes, they are not suitable for the large number of closely related genomes in this work since they require relatively few edges to define a cluster. For example, the MCL algorithm and connected components method assign a node to a cluster as long as it has a single edge, and triangle clustering only requires two edges to two adjacent nodes.

However, connected components and triangle clustering are special cases of the more general *k*-clique percolation algorithm where *k* equals two and three, respectively. The clique percolation algorithm detects clusters by first identifying cliques, sets of nodes which are fully connected, of a specified size *k* in the graph (Fig. 3A). Clusters are then taken as the connected components of an overlap graph where an edge exists between two cliques if they share *k*-1 nodes in common. An intuitive way to visualize this algorithm is by “rolling” a clique of some size *k* over the graph (Fig. 3B). More specifically, a cluster is initiated when a set of nodes which form a *k*-clique is identified. The cluster expands by shifting the *k*-clique to an adjacent *k*-clique that shares *k*-1 nodes in common with the current *k*-clique. A cluster stops expanding when there are no adjacent *k*-cliques, and the algorithm terminates when there are no *k*-cliques which are not part of a cluster. The strength of this algorithm is its ability to exclude sparsely connected nodes from clusters with an easily tunable parameter *k*. Higher values of *k* require greater overlap between a candidate node and those already in the cluster and therefore produce tighter clusters at the cost of excluding more speculative orthology relationships (Fig. 3C). We set *k* to four as compromise between these concerns, yielding 22,813 orthologous groups, a plurality of which contained all 33 species (Fig. S2).

**Fig 3.**
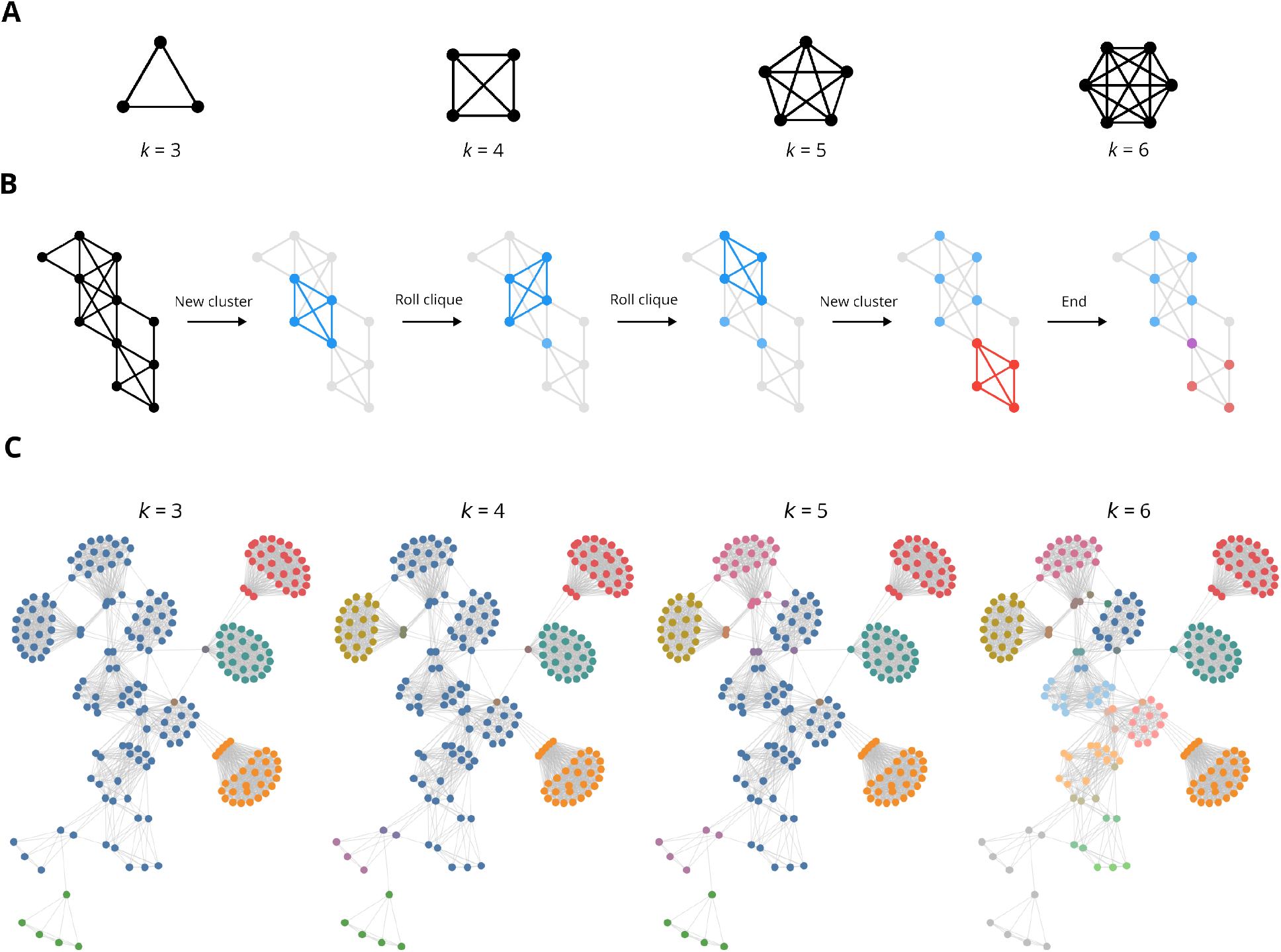
Clique percolation algorithm. **(A)** Cliques for *k* equal to three, four, five, and six. *k* equal to one and two correspond to a single node and two nodes joined by an edge, respectively. **(B)** Illustration of clique percolation algorithm where *k* = 4. **(C)** A single component clustered by clique percolation with varying values of *k*. Nodes are colored according to their cluster. If a node belongs to multiple clusters, it uses a blend of those colors.

### Addition of paralogs to orthologous groups

A weakness of the best reciprocal hits criterion is its exclusion of recently diverged paralogs. Since only the highest scoring hits for each query are included in the graph, a paralog without a corresponding duplicate in the target genome is ignored if it is marginally more diverged than the other copy. This is corrected by adding likely paralogs to the orthologous groups. Briefly, the protein sequences in each genome were searched against themselves. If the bit score for an intra-genome hit exceeded the bit score of any inter-genome hits for the same query, the two sequences were identified as a paralogous pair. Orthologous groups were then supplemented with paralogs by adding the paired sequences for each of the original members of the orthologous group. Most orthologous groups contain no paralogs, and those that do generally have few relative to the original number of sequences in the group (Fig. S3).

### Grouping orthologous groups by gene

As genes can have several annotated isoforms, each gene can be associated with several orthologous groups. However, the orthologous groups are not organized into gene-level units since they were clustered using sequence similarity only. A graph-based approach was therefore used to group orthologous groups with similar sets of parent genes. First, a gene overlap graph was constructed by defining an edge between orthologous groups if the intersection of their associated sets of parent genes is at least 50% of the smaller of the two. Gene groups were then taken as the connected components of the resulting graph, yielding 14,909 groups. This is commensurate with the roughly 15,000 genes in each genome, which suggests this approach has successfully clustered orthologous groups derived from a common set of parent genes.

### Initial alignment and selection of representative sequences

Since the NCBI annotation pipeline incorporates transcriptome data from a variety of sources, its inputs are heterogeneous in sequencing depth, developmental stage, and tissue of origin across different genomes. As a result, some genomes are annotated with different or multiple splice isoforms of a given orthologous gene, which can create complex networks in the resulting hit graph. For example, if the genomes are variably annotated with one or both of two distinct isoforms, the resulting graph may contain two clusters connected by a “bridge” formed by the genomes which contain only one of the isoforms. If the nodes bridging the two clusters form cliques with themselves and the clusters, the clique percolation algorithm will merge all the nodes into a single orthologous group where some genes have multiple associated sequences. However, these additional sequences can complicate downstream comparative analyses that may not easily generalize to genes with multiple associated sequences. Thus, in our pipeline a single representative was chosen for each gene using an alignment-based strategy detailed in the methods section. Briefly, a statistical profile was created from an alignment of the sequences in each orthologous group, and the representative for each gene was chosen as the sequence which best matched this profile.

### Selection of single copy orthologous groups

The criteria for selecting orthologous groups for further analyses depends on the biological question under investigation. For example, studies of gene duplication will focus on orthologous groups with paralogs in some lineages but not in others. In contrast, analyses which assume functional conservation should restrict the orthologous groups to single copy orthologs since paralogs more frequently undergo functional divergence [3, 52, 57]. A simple method for identifying such groups is requiring each species to have exactly one associated gene. However, since the probability of at least one missing gene annotation approaches one as the total number of genomes increases, this is too restrictive and fails to leverage the redundancy of closely related genomes. Instead, a set of phylogenetic diversity criteria detailed in Table S2 were applied to ensure the major lineages were represented in downstream analyses. Furthermore, genome-wide analyses should select one orthologous group per each of the previously identified gene groups as to not bias the results towards genes with many distinct groups of isoforms. In summary, orthologous groups failing the phylogenetic diversity criteria were first removed, and the representative for each gene group was chosen as the highest scoring orthologous group when ranked by the number species and the sum of the bit scores associated with each edge. This significantly reduced the number of orthologous groups from 22,813 to 8,566.

### Alignment refinement

Though the pipeline’s quality control measures ensure a high degree of overall sequence identity between members of an orthologous group, some sequences contain long “poorly supported” segments which have no homology to most or any other sequences in the alignment. Since most common multiple sequence alignment algorithms assume the sequences are largely homologous, these segments are sometimes “over-aligned” by forcing them into alignment where chance sequence similarities occur. Typically, these segments remain contiguous, so the alignments alternate between short runs of columns with few or no gaps and large gap-rich regions (Fig. 4A-B, left). More rarely, when long poorly supported segments are adjacent to a long gap in the same sequence, the two are interlaced, yielding long gaps interrupted by short segments of spurious alignment (Fig. 4C, left).

**Fig 4.**
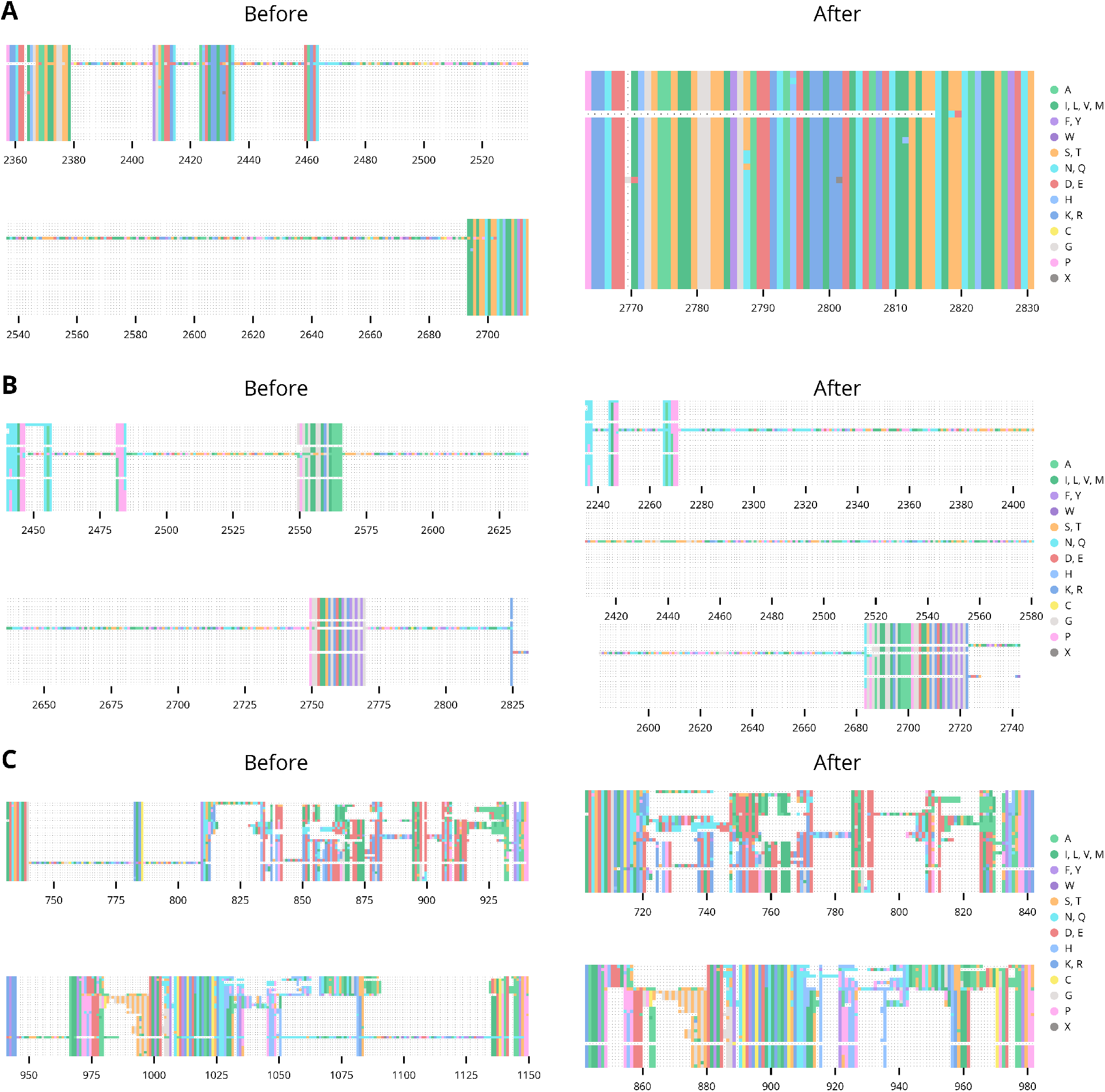
Alignment with long poorly supported segments. The alignments of representative sequences in orthologous groups 0167 **(A)**, 2770 **(B)**, and 23D9 **(C)** before and after refinement.

The aligner MAFFT has a mode for addressing over-alignment with a parameter, *a_max_*, that adjusts the strength of the correction [32, 33]. *a_max_* varies between 0 and 1, with higher values yielding a stronger correction. While values above 0.8 completely eliminate over-alignment and successfully align highly conserved regions, the alignment as a whole is severely degraded, as even homologous sequences with a small amount of divergence are separated by gaps. Thus, in our pipeline orthologous groups were aligned in two stages. In the first, the sequences were aligned with a strong correction of 0.7. Highly conserved regions were identified, which divided the alignment into a complementary set of diverged regions. The sequences in each of these regions were then extracted and aligned separately with a more conservative value for *a_max_* of 0.4. The resulting “sub-alignments” were “stitched” back into their positions in the original alignment. By defining highly conserved “anchor” regions, this approach largely prevents the alignment of chance sequence similarities in long poorly supported segments (Fig. 4, right).

### Alignment curation

Although the refinement process corrects most cases of over-alignment, the alignment may still contain regions whose aligned segments have poor or inconsistent support. For example, long poorly supported segments in internal regions were not removed from the alignment since they are bounded by at least one consensus column to the left and right. Additionally, some regions have a significant fraction of sequences with strongly supported segments, but the observed gap pattern is discordant with the expected phylogenetic relationships. Since they are present in so few sequences, the former segments are likely artifactual, resulting from errors during assembly or annotation. (Biological explanations such as alternative splice sites, frameshift mutations, or transposition events are also possible, however.) In contrast, the high sequence identity and clear boundaries of the segments in the latter regions suggest they are conserved but skipped exons. Given the heterogeneous sourcing of the transcript evidence, these sequences containing these segments are likely splice isoforms specific to certain tissues or developmental stage.

Since the segments in these regions are likely the result of incorrect or incomplete annotations rather than meaningful biological variation, maintaining them in the alignments would propagate spurious homologies to subsequent analyses. This is a common issue in alignments generated by automated pipelines, so downstream analyses often focus on the strongly supported regions by removing or “trimming” columns below some threshold number of gaps or sequence identity [7, 6]. This approach, however, is inadequate if the taxonomic sampling is dense, as a single indel event along a lineage containing many species can increase the number of gaps above the threshold. Moreover, as this method does not incorporate any spatial information, it can rapidly alternate between trimming and preserving columns. Thus, it can severely disrupt any analyses which are sensitive to the spatial organization of an alignment.

Phylogenetic HMMs (phylo-HMMs) are statistical models that incorporate phylogenetic and spatial information to calculate the probability that each observation in a sequence was generated by one of several “hidden states” [18]. Since they can evaluate both the probability of a gap pattern in a column given the known phylogenetic relationships and the local context, a phylo-HMM was used to segment the alignment into contiguous regions with different patterns of gaps. A fully specified phylo-HMM requires a fixed number of hidden states and a probability distribution for each. Thus, we identified four distinct types of regions in the alignments, roughly corresponding to highly conserved regions with few to no gaps, diverged regions, regions with a stable gap pattern discordant with the expected phylogenetic relationships, and regions with poorly supported segments. For simplicity, however, we refer to the states that generate each type of region as 1A, 1B, 2, and 3, respectively. To model probability distributions for each state, we first conceptualized the observed alignments as the superposition of two distinct processes (Fig. 5A, left). The first is a phylogenetic process which evolves and splits a single ancestral sequence over time according to a tree. The second is the annotation process which can erroneously exclude or include segments from a sequence. The result is an alignment of annotated sequences which contains evolutionary information obscured by “noise” from the annotation process, shown here by the exclusion of three N-terminal residues in the fourth annotated sequence. To simplify modeling this behavior with an HMM, we coded the sequences into binary symbols. The distributions for each state then consisted of two components derived from the encoded sequences (Fig. 5A, right). The first component models the gap pattern with a Markov process. This Markov process is in turn composed of two subprocesses where the first is a phylogenetic process, and the second is a jump process. These subprocess roughly correspond to changes caused by evolution and annotation, respectively. Because this first component did not fully capture the propensity for the gap patterns to remain constant, we included a second component that models the “gap stickiness” as a beta-binomial random variable by counting the number of symbols that remain constant between columns. Each component is associated with a set of parameters, and the unique parameters for each state yield its characteristic gap pattern and gap stickiness.

**Fig 5.**
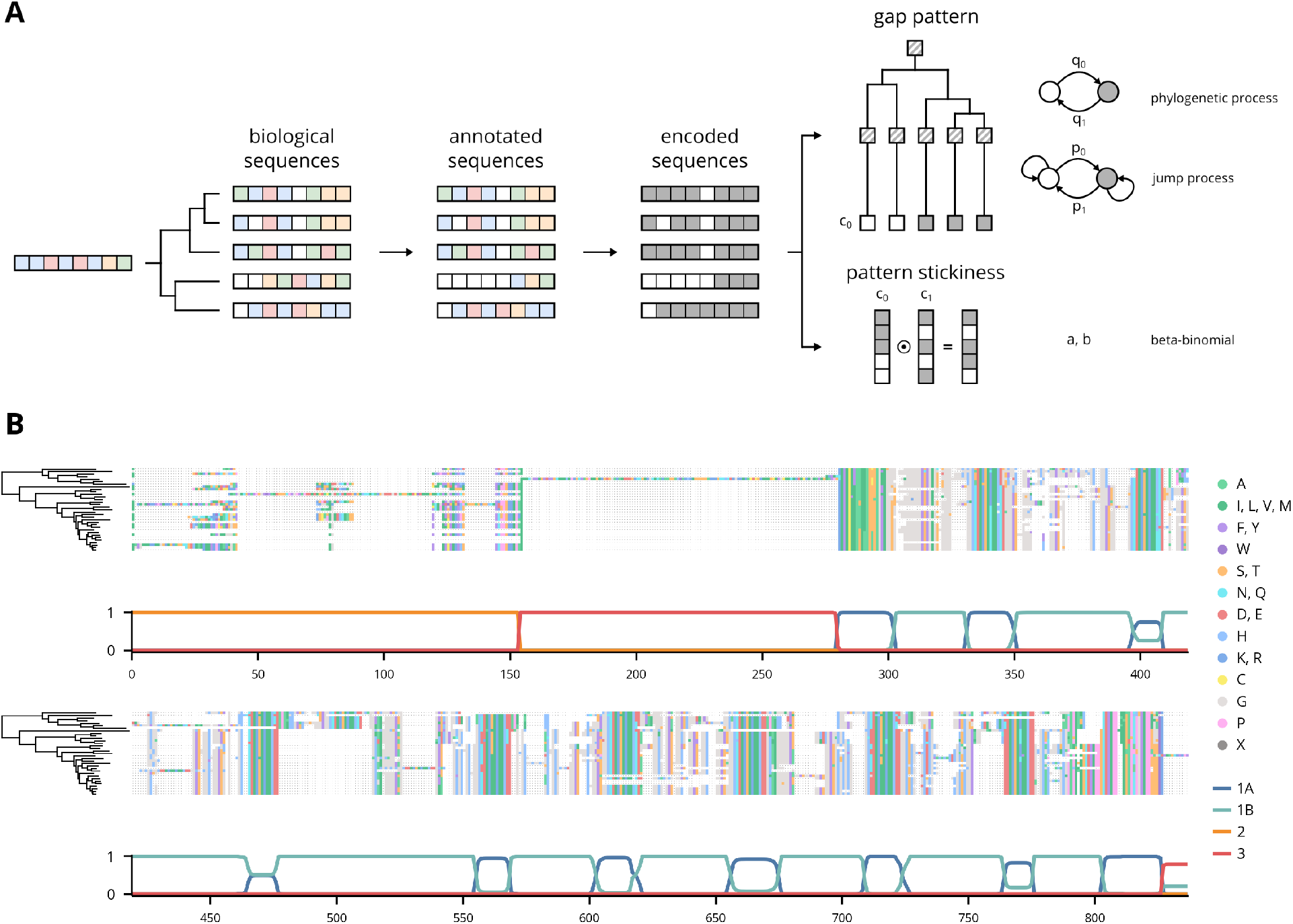
HMM emission architecture and a decoded alignment. **(A)** Schematic of theoretical alignment generating process and corresponding probabilistic components in HMM. White and colored boxes indicate gap and non-gap symbols in the biological sequences, respectively. White and grey boxes indicate gap and non-gap symbols in the encoded sequences, respectively. c_0_ and c_1_ indicate the first and second columns in the alignment, respectively. The parameters associated with each component are shown to the right. **(B)** The alignment of the representative sequences in orthologous group 2252 decoded using the trained HMM.

After the model was trained on manually labeled examples, it was used to assign a label to the columns in each alignment. Columns assigned to states 1A and 1B are the regions of interest for downstream analyses since the gaps generally follow the expected pattern given the phylogenetic tree. In contrast, columns assigned to states 2 and 3 largely corresponded to the long poorly supported segments and phylogenetically discordant regions discussed previously and were therefore removed from the alignments. In the example decoded alignment shown, the decoded states closely follow the expected patterns (Fig. 5B). Overall, 29% of alignments were trimmed of at least one segment or region. However, 87% of the trims were segment trims, meaning the removed segments were largely inferred as state 3 and therefore were likely long poorly supported segments aligned to few if any other sequences (Fig. S5).

Though the phylo-HMM removed phylogenetically incongruent “insertions,” some sequences still contained extensive segments of uninterrupted gaps. These segments are easily identified in regions which are otherwise highly conserved, so they are also likely the result of incorrect or incomplete annotations. However, they can also span more diverged regions, which complicates a simple rule- or threshold-based approach for identifying them. Thus, another phylo-HMM was trained to label each position in a sequence as generated by either a “missing” or “not missing” state. In this case, though, the aligned sequences were processed individually and not as aligned columns. As the previous phylo-HMM already ensured each column has sufficient support, these labels can instead be used to exclude sequences from downstream analyses depending on the amount of tolerated overlap with the regions of interest. Overall, 15% of alignments have at least one sequence with a segment of “missing” data (Fig. S7).

### Inference of species trees

Many phylogenetic methods require a species tree to inform the evolutionary relationships between sequences. In fact, the phylo-HMMs discussed previously used a species tree as an input, though we omitted this detail for clarity of exposition. Therefore, to support the curation step and other downstream analyses, we sought to infer phylogenetic trees from the aligned sequences. However, since the roots of phylogenetic trees are not identifiable with commonly used time-reversible substitution models, we repeated the orthology inference pipeline with the outgroup species *Scaptodrosophila lebanonensis*. Afterwards, we inferred phylogenetic trees using the LG model of amino acid substitution from 100 meta-alignments sampled from alignments of single copy orthologous groups. We then combined them into a single consensus tree (Fig. 6A). To provide a similar tree for the analysis of non-coding regions, we inferred phylogenetic trees using the GTR model of nucleotide substitution from 100 meta-alignments sampled from nucleotide alignments which were “reverse translated” from the protein alignments and their corresponding coding sequences (6B). Both trees have an identical topology, which is consistent with other published phylogenies [9, 36].

**Fig 6.**
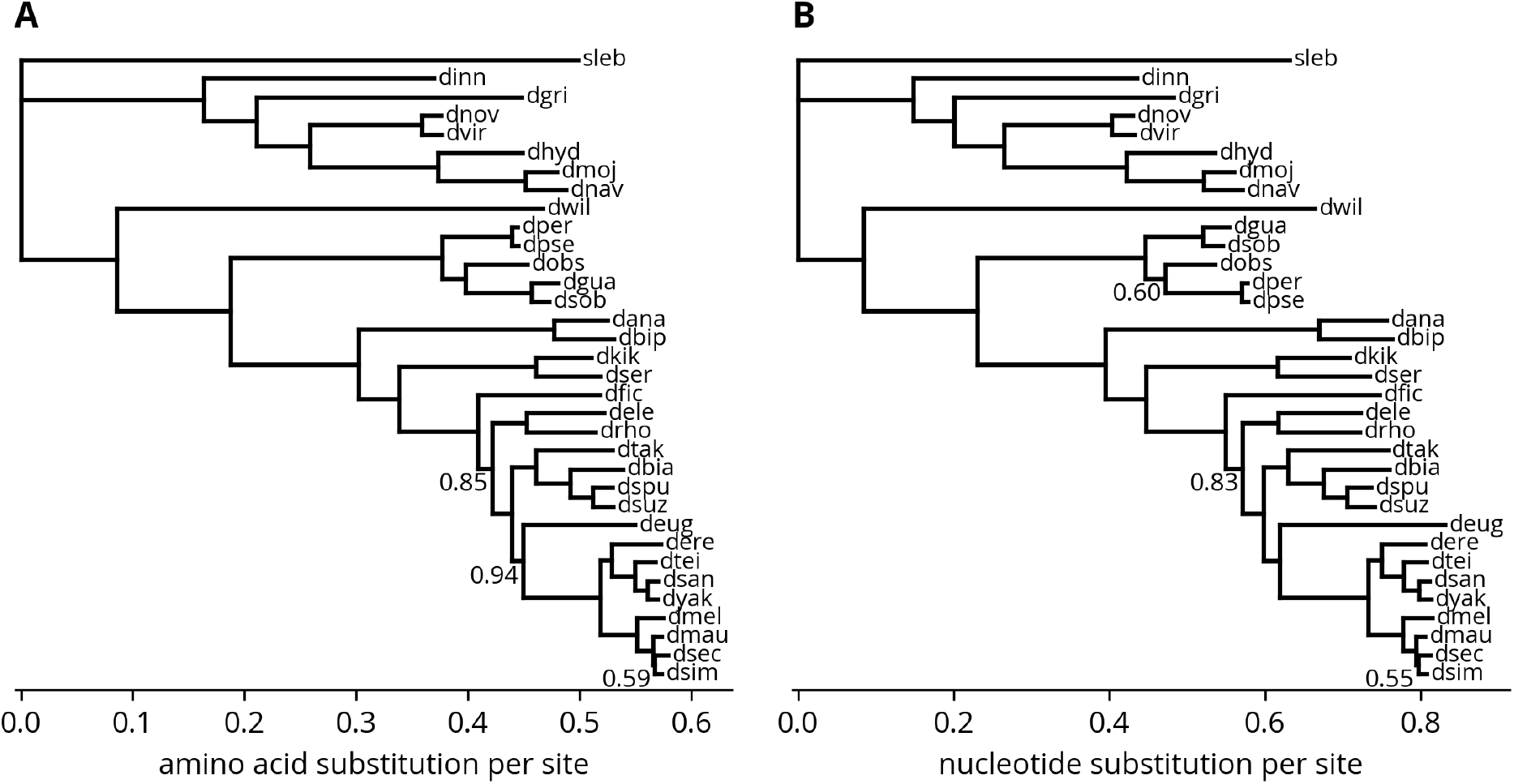
Phylogenetic tree of species. **(A)** Consensus tree from LG model fit to meta-alignments directly sampled from the original protein alignments. **(B)** Consensus tree from GTR model fit to meta-alignments sampled from “reverse translated” nucleotide alignments. Values at nodes are bootstrap percentages.

## Discussion

The orthologous groups and alignments yielded by this pipeline are a valuable resource for comparative studies of gene birth/death processes and protein evolution at the level of both entire proteomes and specific gene families in the *Drosophila* genus. To assist these efforts, the final alignments after curation and the labels from the “missing” phylo-HMM are provided as supplemental data. Although this work focused on single copy orthologs, other studies may require different subsets of orthologous groups that demand other pre-processing and alignment strategies. Therefore, we have included the orthologous groups and the initial alignments with and without non-representative sequences in the supplemental data as well. While we anticipate these resources will remain relevant in the near term, the trends that permitted this work to substantially improve on previous efforts will render them obsolete in the coming years as more *Drosophila* genomes are assembled and annotated. However, an authoritative and lasting set of orthologous groups and alignments is not the primary goal of this work. Instead, it serves as a case study in how dense taxonomic sampling and modern genome assembly and annotation pipelines present new opportunities and challenges to the traditional techniques for identifying and aligning orthologous groups.

For example, despite many additional pre-processing steps and other tweaks introduced by later authors, the basic framework of orthology inference by clustering the hit graph has remained largely unchanged in the past twenty years [58, 53, 38, 30, 40, 16, 61, 10]. This longevity is a testament to the robustness of the underlying idea that orthologous proteins should consistently identify each other as the most similar pairs between their genomes. Even as the number of genomes and their taxonomic density has increased dramatically, many orthology inference pipelines continue to use algorithms which were originally applied to sets of far fewer and more distantly related genomes. This mismatch in scale increases the chance of propagating annotation errors since only a small number of edges are needed to create or merge clusters. Thus, we instead applied a generalization of the triangle and connected components clustering methods called *k*-clique percolation where *k* is a tunable parameter that influences the tightness of a cluster. The optimal value of *k* for a given set of genomes is unclear and likely depends on the desired trade-off between sensitivity and specificity. Furthermore, *k* is not necessarily a global parameter and can instead depend on the properties of each connected component. For example, one possibility is to take an entire component as an orthologous group if its number of unique genes and unique species are equal since all the sequences are isoforms of a single set of genes. This would effectively set *k* equal to one for this component. Another approach is to make *k* an decreasing function of the density of edges, so sparser graphs are clustered more permissively. However, percolation theory or simulations may yield additional insights.

Another challenge is the annotation of multiple isoforms for a single gene. Though prior pipelines have generally selected the longest isoform as the representative before conducting the orthology searches, if the sequences do not share a common intron-exon structure this approach can introduce artifacts or other issues during alignment. Instead, as protein sequences are increasingly derived from or linked to genomic sequences, we sought to incorporate the full annotations into the orthology inference pipeline. This, however, created two additional complications. First, a single gene could have several associated orthologous groups if its isoforms belonged to different clusters. Second, a single orthologous group could have several isoforms of a single gene if its isoforms were clustered together. In both cases, the presence of multiple sequences for a single gene creates ambiguities over which is the “primary” isoform. Since the first occurs at the level of orthologous groups, the orthologous groups were first grouped by the similarity of their parent genes using a graphbased strategy. Afterwards, a single representative is easily chosen as the group with the largest number of distinct species, though other criteria are possible. Since the second occurs within an orthologous group, the sequences were first aligned, and a representative for each gene was chosen as the sequence which was most concordant with this initial alignment.

The second major innovation in this work is its method for the refinement and trimming of alignments. Since sequences produced from automated annotation pipelines can contain long segments which are not homologous to most or any other sequences in their respective orthologous groups, their alignments may contain over-aligned or poorly supported regions which can introduce artifacts into downstream analyses. Thus, in refinement over-alignment is avoided by aligning the sequences in two stages. In the first highly conserved regions are aligned using a strong correction for over-alignment, and in the second more diverged regions are aligned with a weaker correction. This process usually prevents errors caused by long poorly supported segments without degrading the quality of the alignment. In trimming, a phylo-HMM is used to remove regions which are poorly supported by the phylogenetic consensus.

While the combination of these two steps yielded high-quality alignments that are suitable for further analyses, they are an *ad hoc* fix for underlying issues with the gene models and alignment algorithms. The most principled solution is to optimize or supplement the gene models using the initial alignments generated by the orthology inference pipeline, which is possible with tools such as OMGene or OrthoFiller [12, 13]. However, if preserving the original annotations is desired or necessary, the remaining possibility is to correct the alignments as we have done here. In fact, the errors we sought to address broadly stem from shortcomings of current alignment algorithms rather than errors in the sequences themselves. Though the scoring functions of modern multiple sequence alignment algorithms are complex, they are generally derived from models that penalize gaps with a linear or affine cost. As a result, they often interlace gaps with short, aligned segments rather than a single long gap. However, when the sequences are different isoforms of a single gene, their alignment will necessarily contain contiguous exon sized gaps. The same is true when aligning isoforms of diverged orthologs, though the relationship between their exons may be complex.

The most popular aligners for protein sequences (Clustal Omega, MAFFT, MUSCLE, T-Coffee) do not include splice sites in their alignments, which makes them prone to aligning non-homologous exons [56, 32, 14, 48]. Current algorithms can easily be extended to incorporate splice sites by coding them as a new symbol and preventing alignment between splice sites and amino acids, which was recently implemented in the aligner, *Mirage* [47]. The biggest challenge in practice, however, is mapping a protein sequence to its genomic sequence to identify splice sites in the protein sequence. Although this information can in principle be derived from the GTF annotation files produced by the NCBI pipeline, annotating the splice junctions in the protein sequences themselves would facilitate splice-sensitive alignment.

These improvements would enhance rather than replace the phylo-HMM trimming method developed in this work. The model could easily be extended to include a state that outputs a splice symbol before transitioning to one of the states in the current architecture. This intermediate state would increase the accuracy of state inference since a splice symbol followed by a phylogenetically discordant gap pattern would strongly signal a state 2 region. The association between state 2 and skipped exons can be made explicit by requiring that transitions to and from state 2 first proceed through the splice state. This of course depends on proper labeling of the training data, which would be trivial since the boundaries between exons would be marked by splice site symbols rather than inferred from gap patterns. Unfortunately, this would not allow the phylo-HMM to label extended exon boundaries as state 2 since they would not be bounded by splice symbols to the left and right. Accordingly, the phylo-HMM would need to permit transitions between state 3 and any other state to accommodate more complex splice variants and other annotation errors. Thus, this extended phylo-HMM would combine the strengths of splice-sensitive alignment with the more heuristic approach used here. Since state 2 inferences would necessarily correspond to skipped exons, they would be suitable for analyses of this form of alternative splicing. Though state 3 inferences would not directly correspond to specific biological process, they still have value as spatially and phylogenetically aware labels for trimming poorly supported segments from alignments.

The phylo-HMM could be further enhanced by expanding its emission distribution to include more symbols in the amino acid alphabet. This would allow it to better model observed substitution patterns between specific symbols, for example the high rate of exchange between gaps and glutamine residues caused by polyglutamine tracts. The transitions between amino acids could be parametrized with a published matrix such as LG [37], but the transitions between amino acids and gaps would be inferred from labeled data. It is unclear if the resulting gain in accuracy would justify the increased computational burden, however.

Though there are benchmarks available for optimizing and comparing methods of orthology inference, the metrics are calculated over a set of reference genomes which are sparsely sampled over a broad taxonomic range, so it is unclear if they are informative for method designed to yield robust inferences when the genomes are highly related [45]. Furthermore, the heterogeneity of genome architectures and annotations may require quality assurance methods tailored to each set of genomes. Thus, there is likely no one-size-fits-all approach to orthology inference, and with many other standalone programs available for more standard use cases (Hieranoid, OMA standalone, OrthoFinder, OrthoInspector, Orthologer), we have chosen not to package the code into an end-to-end pipeline [31, 2, 15, 39, 65]. Instead, we have devoted considerable attention to organizing and documenting the code to make it accessible to a newcomer and thereby facilitate the adaptation of specific steps to similar projects as needed.

In contrast, though many HMM packages are available for the Python programming language, we found none were satisfactorily documented or contained tutorials to introduce HMMs and their APIs to a wide audience. We therefore refactored this code into a package available on PyPI and GitHub called Homomorph. The package itself only implements standard HMM algorithms, but the GitHub repository includes tutorials that introduce the API and implement training routines. Similar tutorials for machine learning libraries such as TensorFlow have undoubtedly fueled the application of neural networks across diverse fields, but HMMs are also powerful models that can be more appropriate when the data obey certain statistical or structural constraints. Thus, we hope this package and its accompanying tutorials will serve as an on-ramp to HMMs and spur their greater adoption by non-specialists.

Though databases of orthologous groups such as COGs, Ensembl Compara, EggNOG, OMA, OrthoDB, OrthoInspector, and OrthoMCL will continue to be useful for comparative studies across broad taxonomic ranges, the increasing speed at which high-quality genome assemblies and annotations are produced means no single database can encompass the most complete data [23, 27, 28, 1, 65, 44, 8]. Furthermore, since many early comparative genomics studies spanned diverse branches of the tree of life, future research will likely prioritize taxonomic depth over breadth. Thus, custom sets of orthologous groups will grow more and not less common. Despite the challenges these developments pose, they also present new opportunities to bridge the gap between mutational and macroevolutionary processes.

## Materials and methods

### Sequence de-duplication and BLAST search parameters

The protein sequences for each annotation were de-duplicated by removing any sequences which had already appeared in association with the same gene. Thus, the first accession associated with a sequence and gene pair is the sequence’s representative accession for the gene. BLAST+ 2.13.0 was used for the sequence similarity searches [5]. An E-value cutoff of 1 was used for the initial searches. However, this cutoff was lowered to 1E-10 during processing of the BLAST output.

### Extraction of HSPs from BLAST output

To reduce the computational burden of merging HSPs into hits, the BLAST output was filtered to extract HSPs associated with the highest-scoring gene. The HSPs were first grouped by target protein, and the resulting groups were sorted in descending order by the bit score of their highest-scoring HSP. Iterating over the groups, all the HSPs in a group were passed to the next step until the parent gene of the group was not the parent gene of the highest ranked group. This method collects all candidate HSPs for a target gene if the highest-scoring HSP within a group exceeds the highest-scoring HSP of the next best gene. This is in some senses an extension of the best hit criterion where hits are considered at the level of genes rather than proteins. If multiple genes tied for the highest-scoring HSP, the iteration stopped when the parent gene of the current group matched none of these highest-scoring genes.

### Merging of HSPs into hits

HSPs were merged in two stages where the first combined non-overlapping HSPs, and the second combined the remaining HSPs. In the first stage, proceeding from highest to lowest bit score, HSPs were marked as “disjoint” if they do not overlap with any other HSP previously marked as disjoint. Although this greedy strategy does not necessarily yield the highest-scoring set of disjoint HSPs, it prioritized higher scoring HSPs. In the second stage, all disjoint HSPs were marked as “compatible,” and proceeding from highest to lowest bit score the remaining HSPs were marked as compatible if the overlap with any other compatible HSP was no more than 50% of the length of either. The best hit for each query was chosen as the hit with the highest sum of bit scores from disjoint HSPs. The best hits were then filtered by overlap and reciprocity criteria. The overlap criterion was applied first and requires that 50% of residues in the query are aligned in compatible HSPs. This excludes false positives from conserved domains embedded in larger non-homologous proteins by ensuring the hits span a sufficient fraction of the query and target sequences. The reciprocity criterion requires each query-target pair has a corresponding hit where the roles are reversed, which ensures there is no ambiguity in which target is the best match for the query.

### Clustering by *k*-clique percolation

*k*-clique percolation was implemented in two steps. In the first, maximal cliques were identified. In the second, a percolation graph is constructed by defining edges between cliques if they have *k*-1 nodes in common. Clusters are the connected components of this second graph. The first step uses the NetworkX implementation of a maximal clique algorithm. The second step uses a modification of the NetworkX implementation of the *k*-clique community algorithm. The NetworkX implementation exhaustively finds all edges in the percolation graph. Since joining a *k*-clique community only requires that a clique has a single edge connecting it to that a community, this approach is needlessly expensive for large graphs. The custom implementation instead uses a progressive approach where each clique is checked against a list of known communities, merging communities as necessary in each step.

The hit graph is sparse, so these algorithms are efficient when applied to its individual connected components. However, some components have a structure with many maximal cliques, which dramatically slows the first or second step of the clique percolation algorithm. Thus, if either step exceeded 90 s, the process timed out, and the simpler *k*-core algorithm was used instead. Out of over 10,000 connected components, only seven timed out, and many of those contained highly dense clusters of histone sequences.

### Addition of paralogs to orthologous groups

The protein sequences for each annotation were searched against themselves with the same settings as for the inter-genome searches. The resulting output was processed identically except the HSPs were not filtered using the best gene criterion. Thus, all HSPs for each query were merged into hits. The best hit for each query and target gene was then chosen as the hit with the highest sum of bit scores from disjoint HSPs. (Grouping by target gene ensured only the highest-scoring isoform was selected.) Query-target pairs whose hits which exceeded the maximum bit score for all inter-genome hits associated with that query and passed the overlap and reciprocity filters were designated as paralogous pairs. The orthologous groups were supplemented with paralogs by adding the paired sequences for each of the original members of the orthologous group.

### Initial alignment and selection of representative sequences

The sequences in each orthologous group were aligned using MAFFT 7.490 with the following settings: --globalpair --maxiterate 1000 --thread 1 --anysymbol --allowshift --leavegappyregion --unalignlevel 0.4 [32]. Representative sequences for each gene were then selected by maximum likelihood according to binary profiles constructed from these alignments. First each sequence was coded into gap and non-gap symbols. The sequences were then grouped by gene, and for each group and position if at least one sequence was aligned in the group, the group contributed one count for the non-gap symbol to the profile at that position. Otherwise, the group contributed a count for the gap symbol at that position. To account for the phylogenetic dependencies between sequences, the counts were weighted according to a Gaussian process over the GTR2 consensus tree described in the section on inferring species trees [4]. If a species had multiple genes in the orthologous group, the species weight was divided evenly among them. Each coded sequence was then scored according to this profile, and the maximum likelihood sequence for each gene was selected as its representative. By assigning a non-gap count to groups and positions where at least one sequence is aligned, the profile incentivizes the selection of sequences contain the fewest gaps and match the consensus alignment. Unfortunately, this scheme can cause a sequence to score negative infinity if it has a gap at a position where every group has at least one aligned sequence, so the profile was initialized with a pseudocount of 0.005 for the gap and non-gap symbols at each position.

### Alignment refinement

The representative sequences in the single copy orthologous groups were aligned with the same settings as described in the previous section except *a_max_* was set to 0.7. A binary profile was created from the alignment using Gaussian process sequence weighting as described in the section on selecting representative sequences. (Because the orthologous groups were single copy and contained only representative sequences, each sequence received the full weight associated with its species.) The binary profile was then converted into a binary mask by identifying where the weighted fraction of non-gap symbols exceeded 0.5. The binary mask was closed with a structuring element of size three, and highly conserved regions were identified as the contiguous intervals of this closed mask with a minimum length of 10. Diverged regions were taken as the complement of the highly conserved regions. For each diverged region, the corresponding segments of the sequences in the initial alignment were extracted and aligned with *a_max_* set to 0.4. The resulting sub-alignments were then stitched into the initial alignment.

### Alignment curation

The alignments were coded into gap and non-gap symbols to simplify the emission distributions. The “insertion” phylo-HMM was composed of the four hidden states described in the main text. The emission distributions for each consisted of two components which modeled the gap pattern and the propensity for those patterns to remain constant (“gap stickiness”), respectively. The first component was a two-state Markov process which was in turn composed of two subprocess. The first was a phylogenetic process on the on the GTR2 consensus tree described in the section on inferring species trees, and the second was jump process at the tips. The second component was a beta-Bernoulli distribution on the number of symbols which were constant between subsequent columns. The “missing data” phylo-HMM was composed of two hidden states which were both parameterized with the same two-state, two component Markov process as the insertion phylo-HMM. However, the emission probabilities were calculated as the posterior probability of the observed symbol given the data rather than the probability of the data. Only the alignments of the single copy orthologous groups were curated, so each tip in the species tree would correspond to a single sequence in the alignment.

All HMM algorithms were implemented with custom code which is available as the package Homomorph on PyPI. The insertion phylo-HMM was trained on 47,387 manually labeled columns in 14 alignments, and the deletion phylo-HMM was trained on 67,001 manually labeled positions in 23 sequences in 11 unique alignments (Fig. S4, Fig. S6). Because maximum-likelihood estimation of the model parameters yielded posterior decoding curves which toggled between hidden states too rapidly, the models were instead trained discriminatively [35]. The difference, briefly, is maximum-likelihood estimation finds the parameters that best reproduce the observed distributions whereas discriminative training finds the parameters that minimize prediction error. Discriminatively trained models typically perform better in practice since real-world data are rarely fully described by the distribution specified by the model.

The posterior distributions over states were calculated for each alignment using the trained insertion phylo-HMM. Regions with a high probability of state 2 or 3 are candidates for trimming. However, because the probability of a state can change rapidly or gradually depending on the local context, a simple cutoff would not necessarily define the boundaries of these regions as the columns where the gap pattern changes most abruptly. Instead, the following algorithm was used. First, a high cutoff defines a “seed” region. The left and right endpoints of the seed are then expanded both inwards and outwards to define two intervals from which boundaries are selected. The outward expansion halts when the probability or its derivative is below two distinct thresholds, respectively. The inward expansion halts when the derivative is below a different threshold. The left and right boundaries are chosen as the columns in each interval with the maximum product between the derivative and the change in the gap profile between columns. The gap profile is calculated as the number of gaps in each column using Gaussian process sequence weighting as described in the section on selecting representative sequences. By combining where the model’s confidence changes rapidly with the observed change in the gap pattern, this method generally selects reasonable boundaries.

States 2 and 3 have distinct characteristics which require different trimming strategies, so regions with a high probability of state 3 were handled first. Because the long poorly supported segments in state 3 regions were sometimes aligned to highly conserved columns or short segments in other sequences, trimming columns entirely would remove these segments from the other sequences even if they do not qualify as long and poorly supported themselves. Thus, regions with high state 3 probabilities were trimmed at the level of individual sequences rather than entire columns with the following method. First, the probability of state 3 of columns with a gap profile value less than or equal to 0.1 was set to 0 to break long poorly supported segments aligned to highly conserved columns into separate regions. Regions were then defined with the previously described algorithm using high and low cutoffs of 0.75 and 0.01, respectively. The outer and inner derivative cutoffs were both 0.001. The mean number of non-gap symbols in a region was calculated using Gaussian process sequence weighting. (The sequences with the five most non-gap symbols were also excluded to not bias the estimate with long poorly supported segments.) The final mean *μ* was taken as the minimum of this value and two. A cutoff *k*, derived from a geometric model of the number of non-gap symbols and a significance level α, was calculated using the following equation *k* = log(*α*)/log(1 – *p*) – 1 where 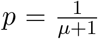 and *α* = 0.01. Any sequence whose number of non-gap symbols in the region equaled or exceeded this value was trimmed by replacing all non-gap symbols with gaps.

To trim the remaining state 2 regions, the posterior probability of state 2 was added to a modified state 3 probability which was set to zero for any state 3 regions identified in the previous step. This ensured that any regions which were intermediate between state 2 and 3 were included. Regions were defined from this combined probability using the algorithm described previously except with a high probability cutoff of 0.9 instead of 0.75. The posterior probabilities from the missing data phylo-HMM were also converted into state assignments using a similar method. However, the initial seeds were defined with a cutoff of 0.75, and the seeds were expanded outward to first nongap symbol or until the posterior probability of the “missing data” state was below 0.05. These assignments, which are available in the supplementary data, can be used to filter segments or entire sequences from downstream analyses.

### Inference of species trees

The orthology inference pipeline was first repeated with the outgroup species *Scaptodrosophila lebanonensis*. Then orthologous groups with one sequence for each species were aligned, and 100 meta-alignments were constructed by randomly sampling 10,000 columns from these 9,435 alignments. (The alignments were not refined before sampling.) To determine the effect of invariant columns and gaps, two sampling strategies were used where invariant columns were allowed or disallowed and the maximum fraction of gaps was set at 0, 50, and 100%. Their combination yielded six different sets of meta-alignments. A tree was fit to each meta-alignment with the LG substitution model and four discrete gamma rate categories using IQ-TREE 1.6.12 [46, 37, 64]. If the sampling strategy allowed invariant columns, an invariant rate category was included. The resulting trees from each set were then merged into a majority consensus tree (Fig. S8).

To fit trees using the GTR model of nucleotide substitution, the protein alignments were converted to nucleotide alignments using the corresponding coding sequences in the genome annotations. Some protein sequences are “low quality,” meaning their coding sequences contain frameshifts, premature stop codons, or other errors even though they are strong hits to known protein-coding genes. The NCBI annotation pipeline corrects some of these defects in the protein sequences, which can complicate a simple “reverse translation” of the alignment. After rejecting alignments where the expected translation from a coding sequence differed from its corresponding protein sequence, 3,425 alignments remained. Consensus tree were derived from meta-alignments sampled from these alignments using the approach described previously except the GTR model was used in place of the LG model.

To fit trees using the two-state GTR model of substitution, the protein alignments were first coded into gap or non-gap symbols. As before, 100 meta-alignments were constructed from these coded alignments for each sampling strategy. In this case, only the presence of invariant columns was varied, yielding two sets of meta-alignments. Trees were then fit using the GTR2 model with no rate categories. An invariant category, however, was included if invariant columns were allowed. Since the bootstrap confidences were sometimes lower than 50%, the resulting trees from each set were then merged into a loose consensus tree to prevent multifurcations (Fig. S9).

## Footnotes

1 For multiple sequences, orthology is usually defined relative to their most recent common ancestor. This technically includes sequences which split by duplication after this point (“in-paralogs”), but excludes sequences which split by duplication before (“out-paralogs”) [53]. Many current orthology inference pipelines explicitly incorporate steps to detect in-paralogs. However, the resulting orthologs groups can easily be restricted to those without in-paralogs (“single copy orthologs”) for analyses where an assumption of conserved function is necessary.

## Code and data availability

The code used to produce the results and analyses is available at https://github.com/marcsingleton/orthology_inference2023. HMM algorithms were implemented in the standalone package Homomorph which is available at https://github.com/marcsingleton/homomorph and on the Python Package Index (PyPI). The following Python libraries were used: matplotlib, NumPy, pandas, SciPy, and TensorFlow [29, 26, 42, 62, 41]. Relevant output files are available in the supporting information. There is no primary data associated with this manuscript. All primary data are available from publicly accessible sources described in their corresponding sections.

## Acknowledgments

We thank the past and present members of the Eisen lab for helpful discussions and their assistance in revising the manuscript.

## Supporting information

**Fig S1.**
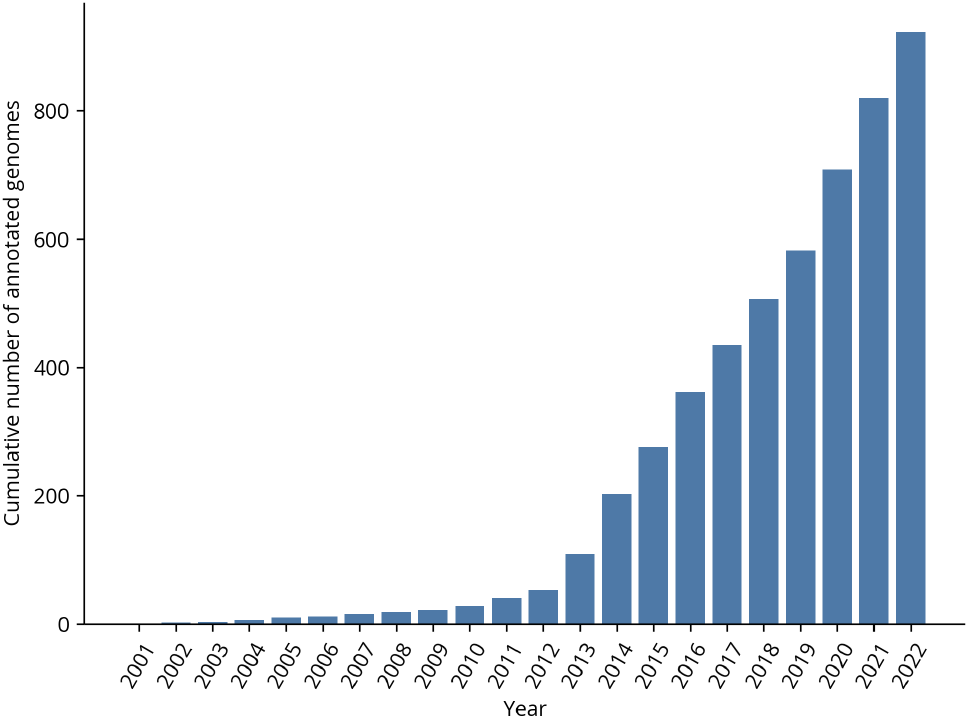
Cumulative number of different eukaryotic genomes annotated by NCBI.

**Fig S2.**
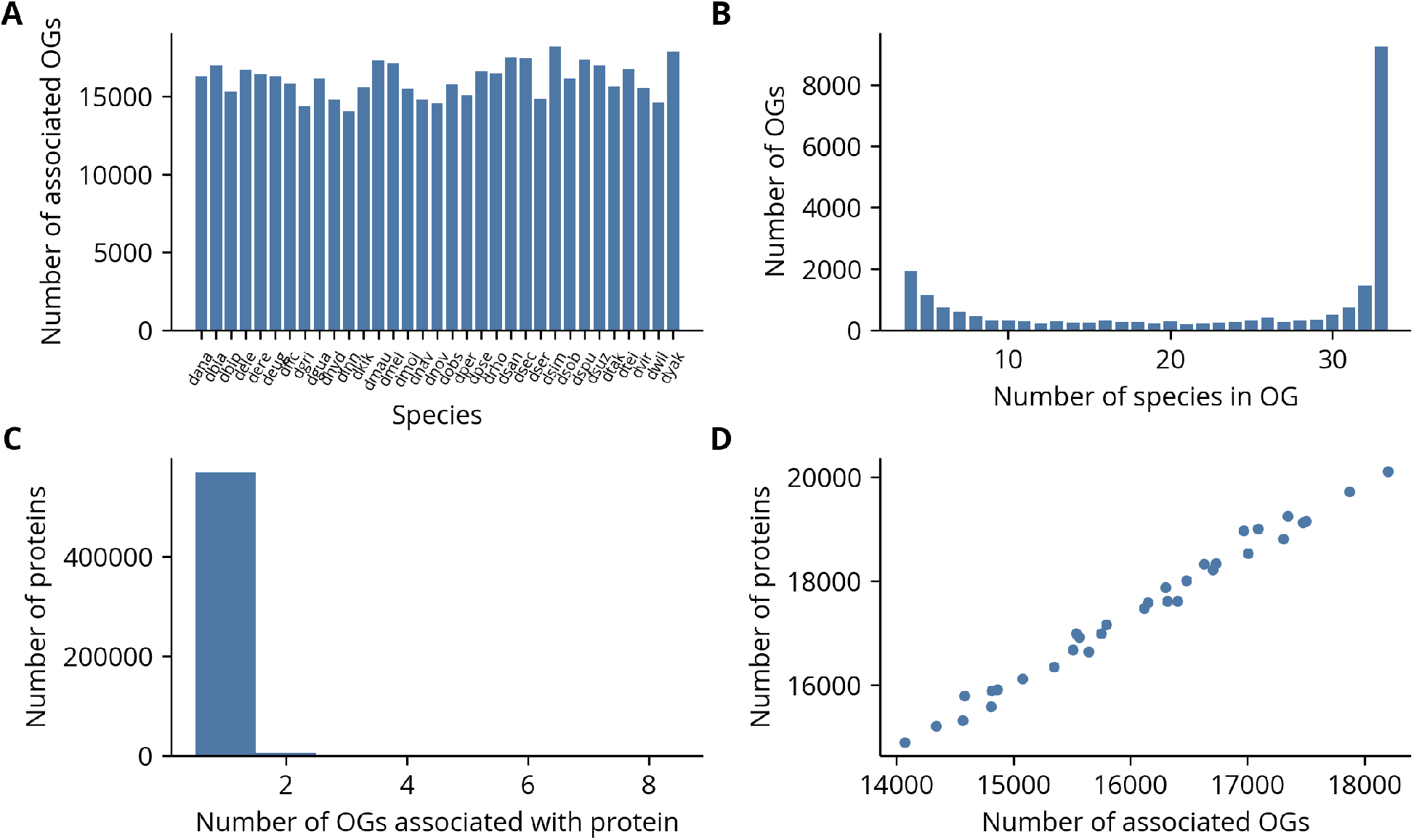
Statistics of orthologous groups. **(A)** Each species is equally represented in orthologous groups (OGs). **(B)** A plurality of orthologous groups contain all species. **(C)** Nearly all proteins are associated with a single orthologous group. **(D)** The number of orthologous groups associated with a species is strongly correlated with the number of unique annotated proteins, which suggests the annotation pipeline generally identifies conserved genes.

**Fig S3.**
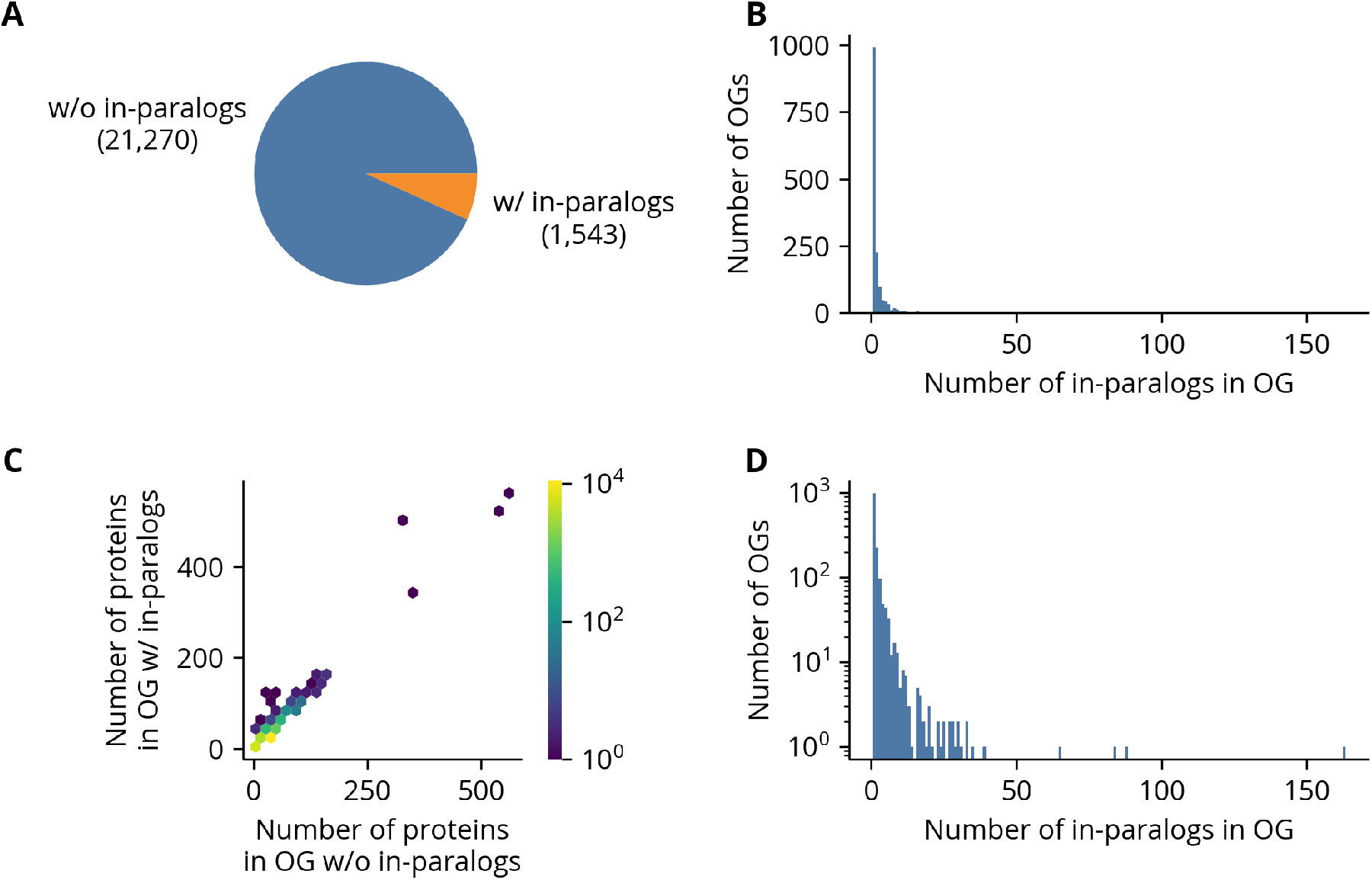
Addition of paralogs to orthologous groups. **(A)** Most orthologous groups (OGs) have no in-paralogs. **(B, D)** Of the groups with paralogs, most have fewer than five. **(C)** The in-paralogs are generally only a small fraction of the sequences in an orthologous group.

**Fig S4.**
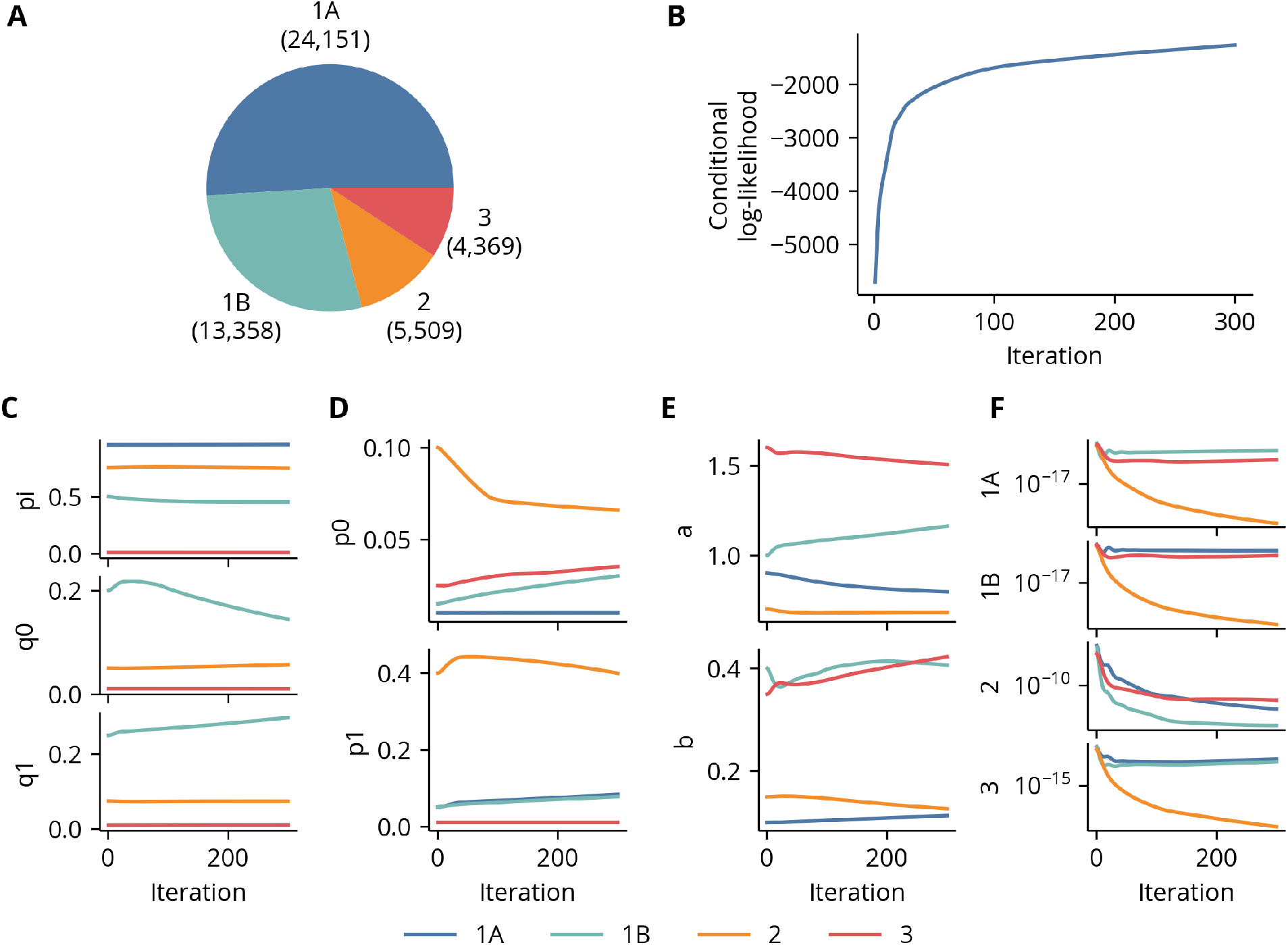
Insertion phylo-HMM data and training details. **(A)** Most columns in the training data were labeled as state 1A or 1B. **(B)** The model loss stabilized by the final training iteration. **(C-F)** The values of parameters in the phylogenetic process, the jump process, the pattern stickiness model, and the transition matrix, respectively, at each training iteration. The transition matrix plots are the transition rates to the state indicated on the vertical axis and given in log scale. Self transitions are excluded.

**Fig S5.**
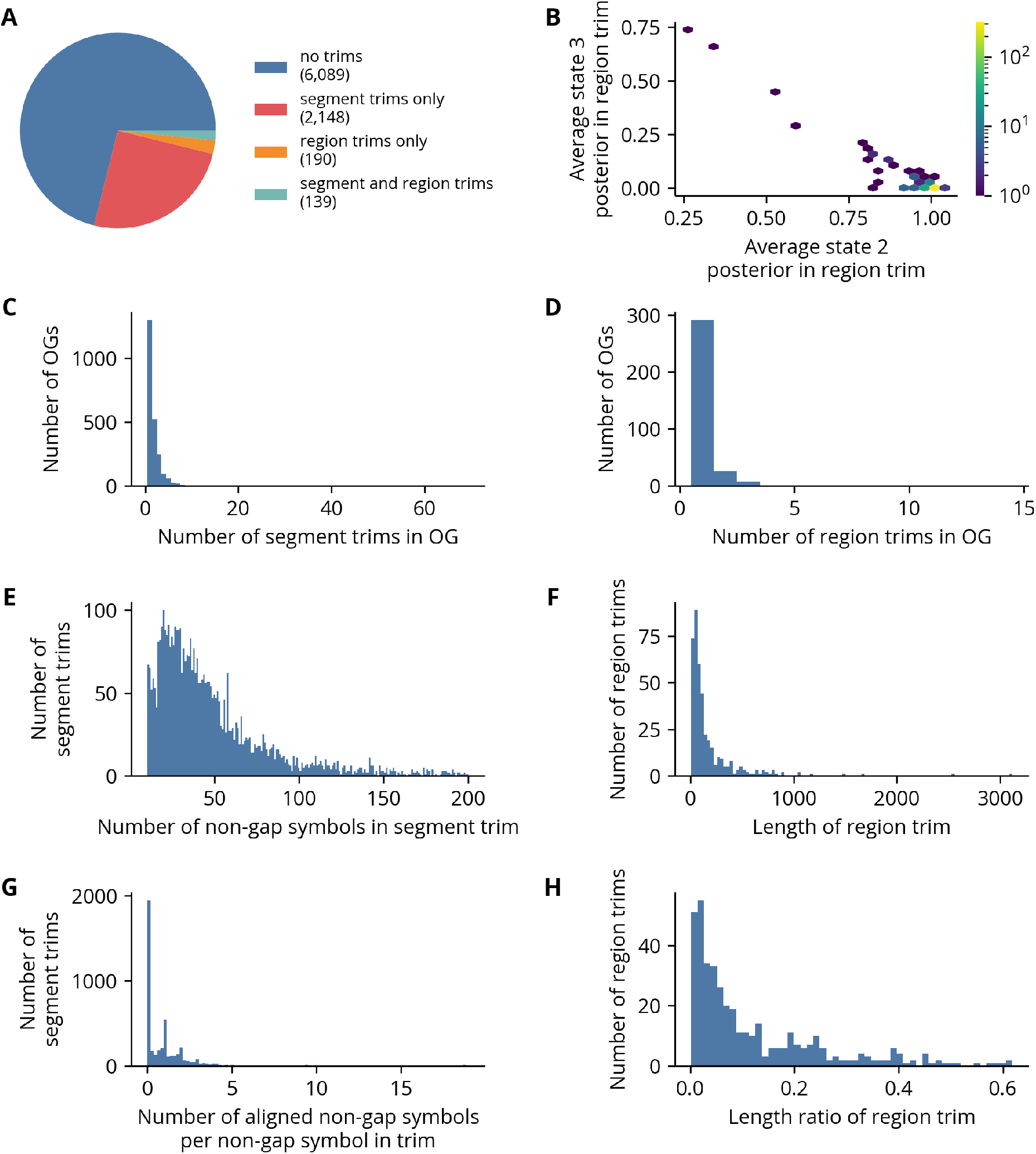
Insertion phylo-HMM trimming details. **(A)** Most alignments were not trimmed. Of the alignments with trims, most were trimmed only at the level of sequences. **(B)** Most trimmed regions were inferred primarily as state 2. **(C)** Most alignments with sequence trims have fewer than 10 segments removed. **(D)** Most alignments with region trims have fewer than five regions removed. **(E, G)** The number of non-gap symbols in sequence trims can vary considerably, but for nearly all sequence trims each non-gap symbol in the removed segment is aligned to fewer than five non-gap symbols on average. Only the lower 95% of the distribution of the number of nongap symbols in the sequence trims is shown. **(F, H)** The length of region trims can also vary considerably, but generally each region trims accounts for fewer than 10% of the columns in the original alignment.

**Fig S6.**
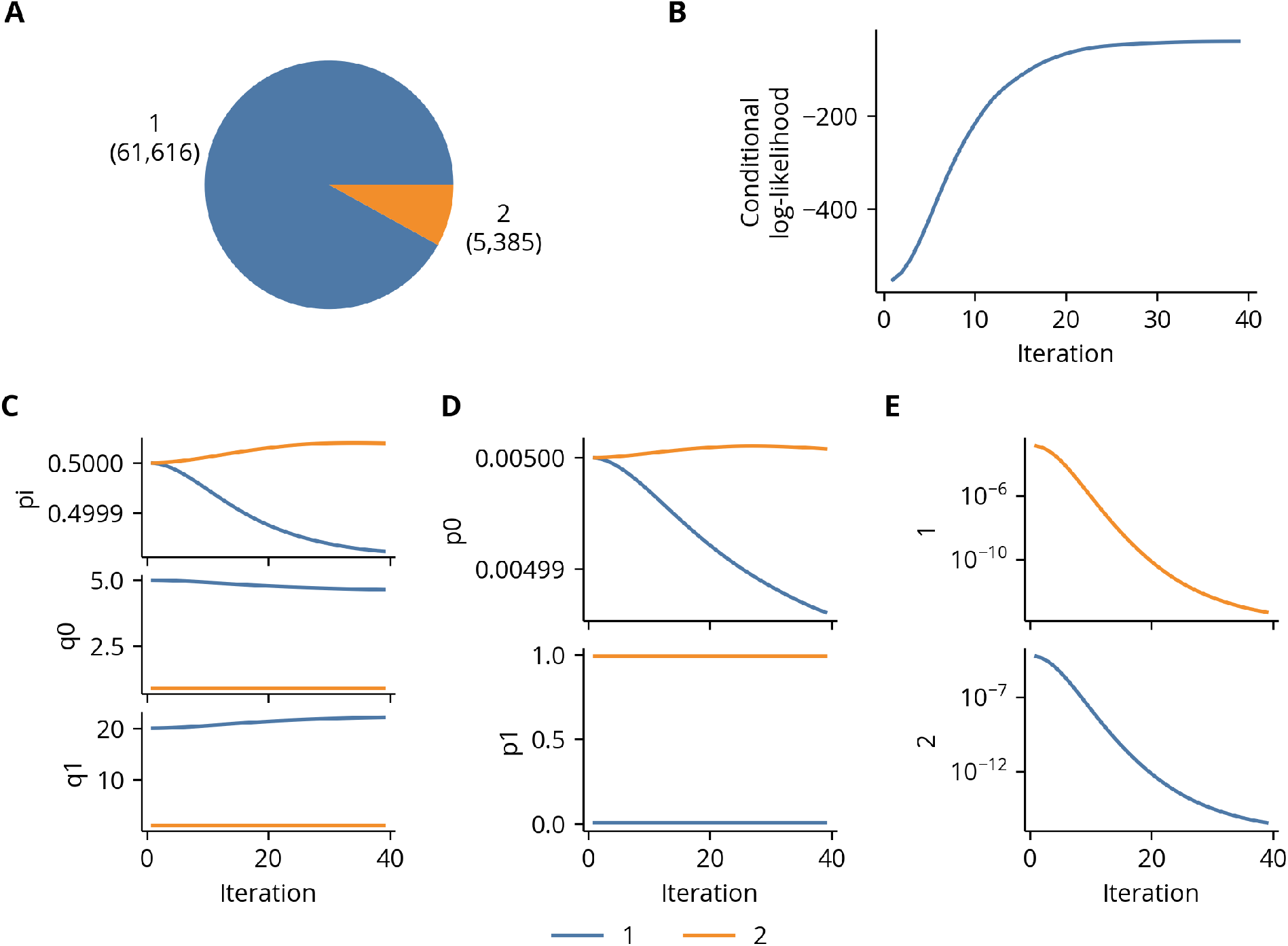
Missing phylo-HMM data and training details. **(A)** Most columns in the training data were labeled as state 1, which is referred to as the “not missing” state in the main text. **(B)** The model loss stabilized by the final training iteration. **(C-F)** The values of parameters in the phylogenetic process, the jump process, and the transition matrix, respectively, at each training iteration. The transition matrix plots are the transition rates to the state indicated on the vertical axis and given in log scale. Self transitions are excluded.

**Fig S7.**
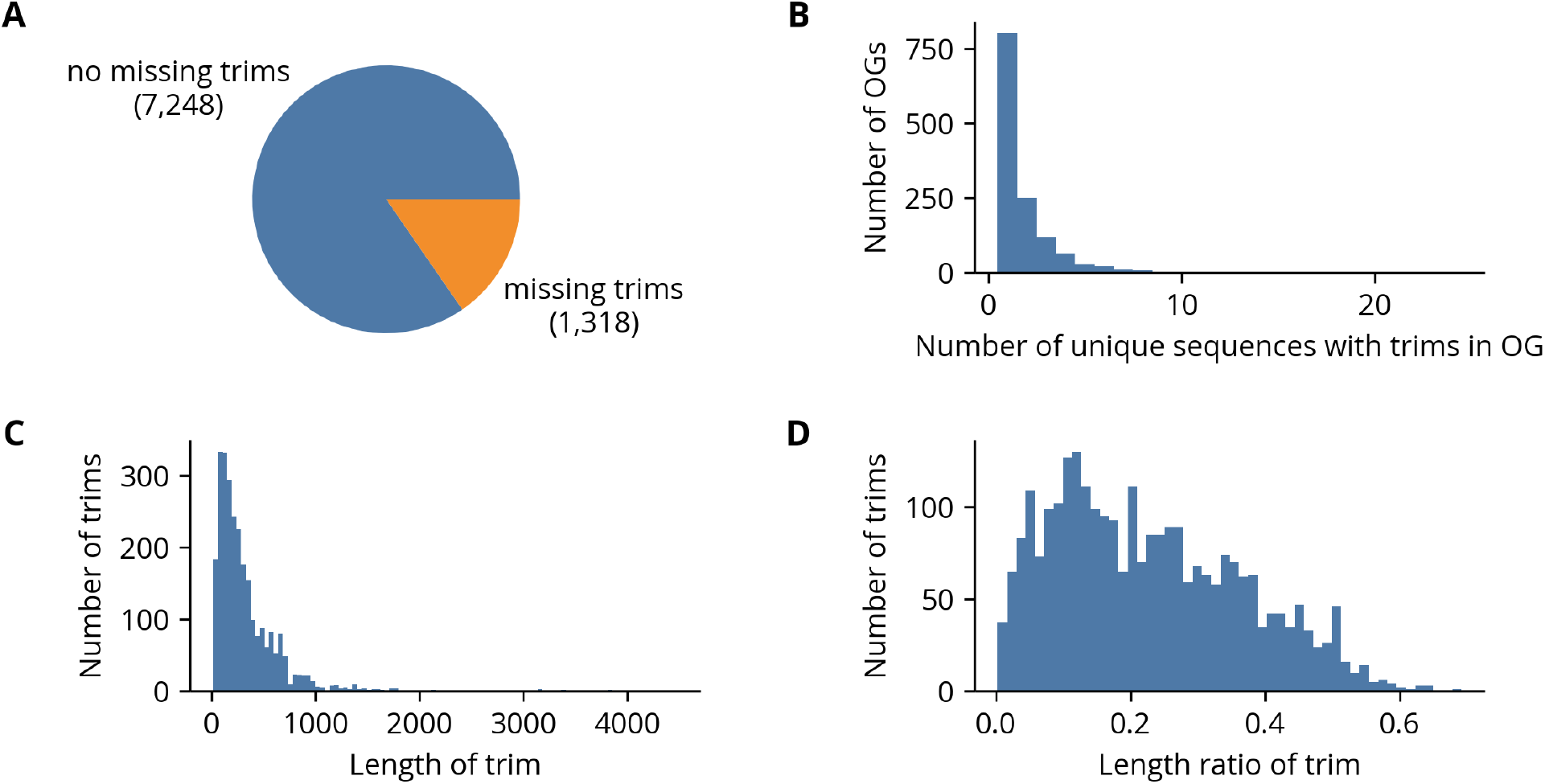
Missing phylo-HMM trimming details. **(A)** Most alignments have no sequences with “missing” segments. **(B)** Of the alignments with sequences trimmed of missing segments, a majority have only one trimmed sequence. **(C-D)** The length of missing segments can vary considerably, both in terms of the number of positions as well as its ratio to the number of columns in the alignment.

**Fig S8.**
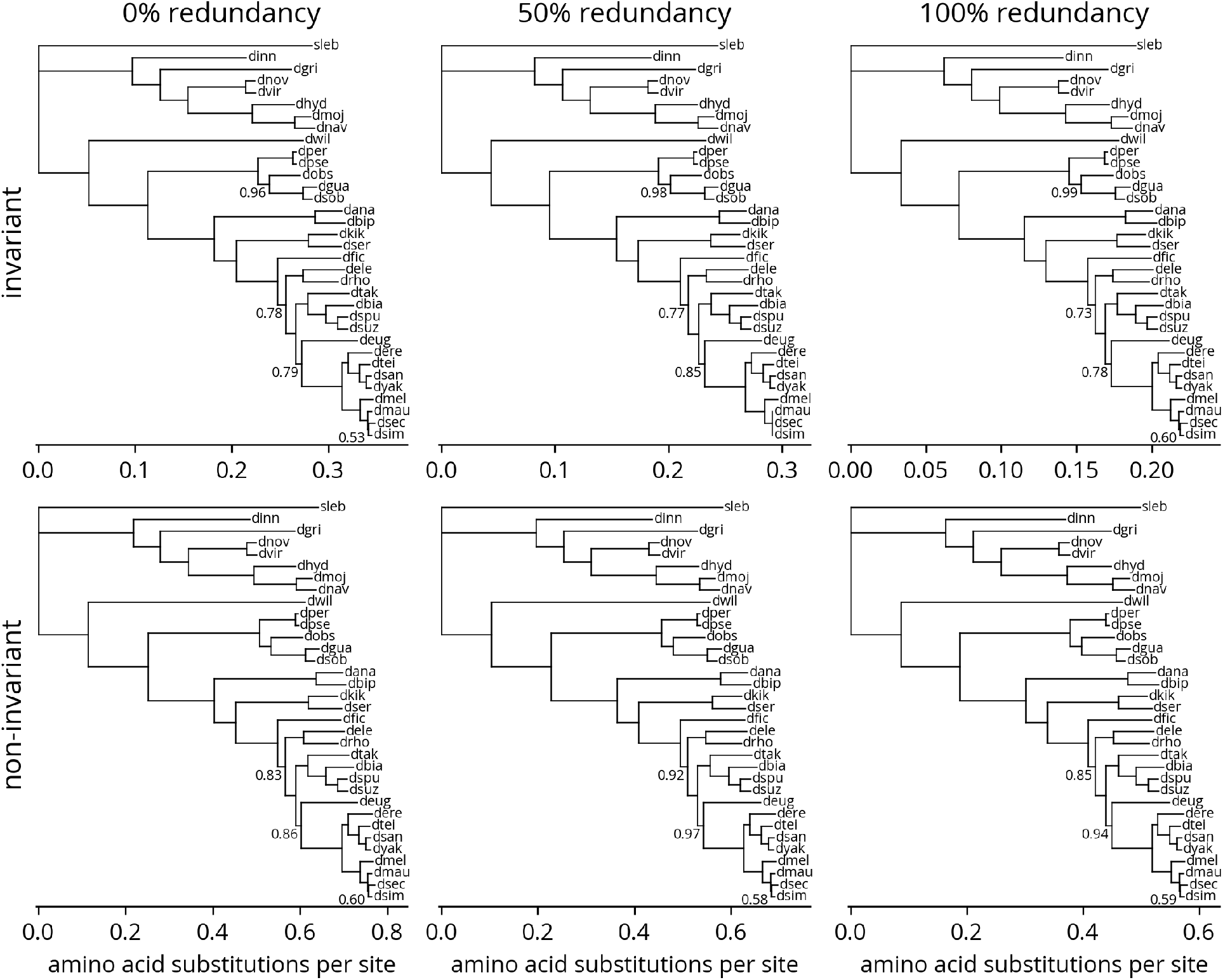
Phylogenetic trees created by different sampling strategies under LG model.

**Fig S9.**
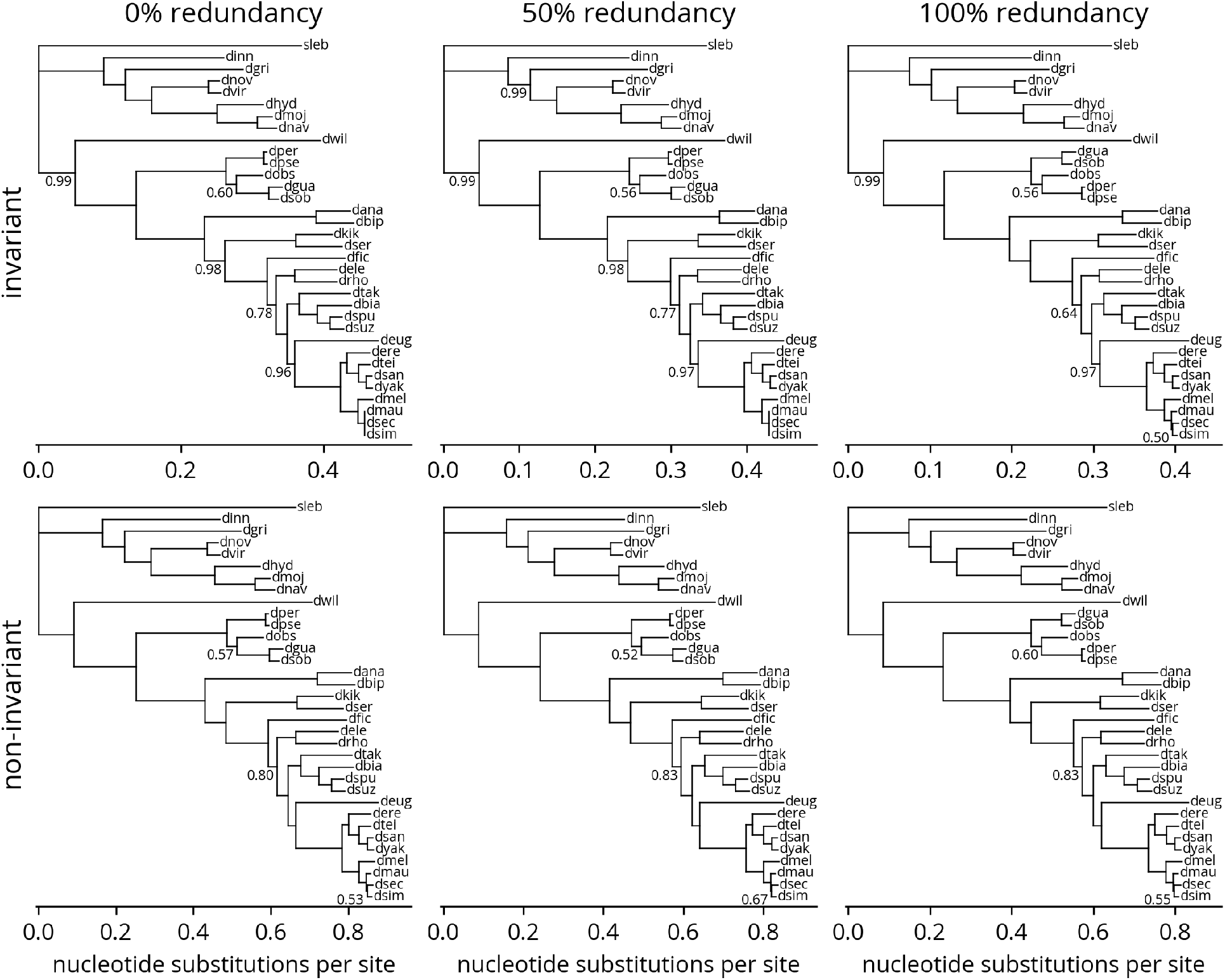
Phylogenetic trees created by different sampling strategies under GTR model.

**Table S1.** Genome annotations.

| **Species** | **Species ID** | **Taxon ID** | **Version** | **Source** |
| --- | --- | --- | --- | --- |
| *Drosophila ananassae* | dana | 7217 | 102 | NCBI |
| *Drosophila biarmipes* | dbia | 125945 | 102 | NCBI |
| *Drosophila bipectinata* | dbip | 42026 | 102 | NCBI |
| *Drosophila elegans* | dele | 30023 | 102 | NCBI |
| *Drosophila erecta* | dere | 7220 | 101 | NCBI |
| *Drosophila eugracilis* | deug | 29029 | 102 | NCBI |
| *Drosophila ficusphila* | dfic | 30025 | 102 | NCBI |
| *Drosophila grimshawi* | dgri | 7222 | 103 | NCBI |
| *Drosophila guanche* | dgua | 7266 | 100 | NCBI |
| *Drosophila hydei* | dhyd | 7224 | 101 | NCBI |
| *Drosophila innubila* | dinn | 198719 | 100 | NCBI |
| *Drosophila kikkawai* | dkik | 30033 | 102 | NCBI |
| *Drosophila mauritiana* | dmau | 7226 | 100 | NCBI |
| *Drosophila melanogaster* | dmel | 7227 | FB2022_02 | FlyBase |
| *Drosophila mojavensis* | dmoj | 7230 | 102 | NCBI |
| *Drosophila navojoa* | dnav | 7232 | 101 | NCBI |
| *Drosophila novamexicana* | dnov | 47314 | 100 | NCBI |
| *Drosophila obscura* | dobs | 7282 | 101 | NCBI |
| *Drosophila persimilis* | dper | 7234 | 101 | NCBI |
| *Drosophila pseudoobscura* | dpse | 7237 | 104 | NCBI |
| *Drosophila rhopaloa* | drho | 1041015 | 102 | NCBI |
| *Drosophila santomea* | dsan | 129105 | 101 | NCBI |
| *Drosophila sechellia* | dsec | 7238 | 101 | NCBI |
| *Drosophila serrata* | dser | 7274 | 100 | NCBI |
| *Drosophila simulans* | dsim | 7240 | 103 | NCBI |
| *Drosophila subobscura* | dsob | 7241 | 100 | NCBI |
| *Drosophila subpulchrella* | dspu | 1486046 | 100 | NCBI |
| *Drosophila suzukii* | dsuz | 28584 | 102 | NCBI |
| *Drosophila takahashii* | dtak | 29030 | 102 | NCBI |
| *Drosophila teissieri* | dtei | 7243 | 100 | NCBI |
| *Drosophila virilis* | dvir | 7244 | 103 | NCBI |
| *Drosophila willistoni* | dwil | 7260 | 102 | NCBI |
| *Drosophila yakuba* | dyak | 7245 | 102 | NCBI |
| *Scaptodrosophila lebanonensis* | sleb | 7225 | 100 | NCBI |

**Table S2.**
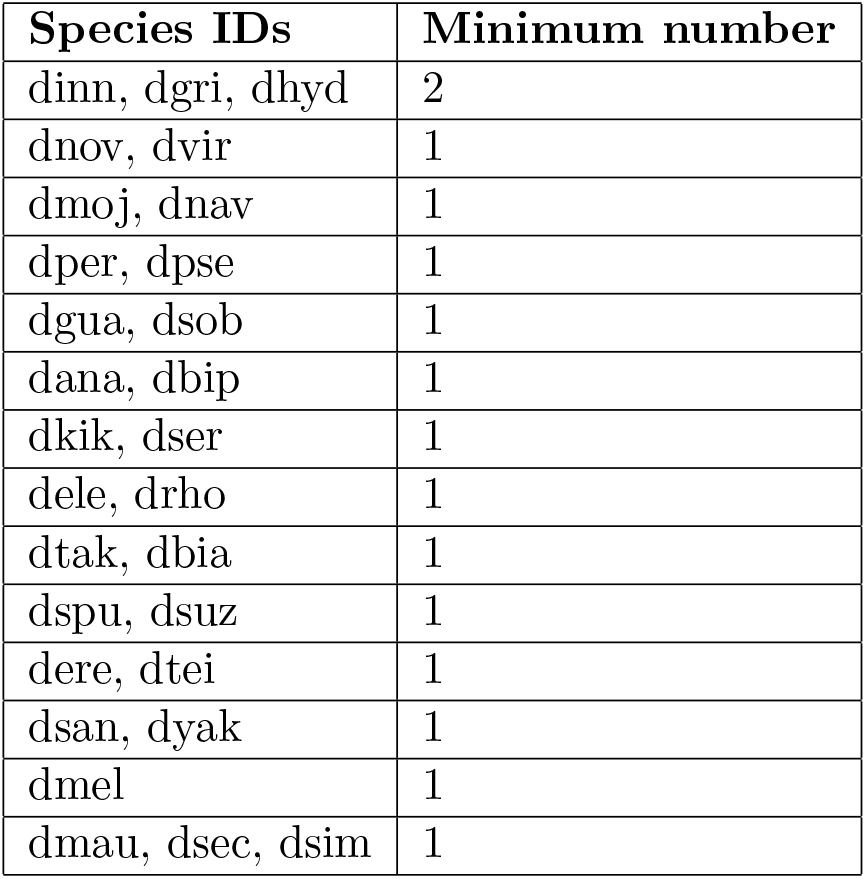
Phylogenetic diversity criteria.

| **Species IDs** | **Minimum number** |
| --- | --- |
| dinn, dgri, dhyd | 2 |
| dnov, dvir | 1 |
| dmoj, dnav | 1 |
| dper, dpse | 1 |
| dgua, dsob | 1 |
| dana, dbip | 1 |
| dkik, dser | 1 |
| dele, drho | 1 |
| dtak, dbia | 1 |
| dspu, dsuz | 1 |
| dere, dtei | 1 |
| dsan, dyak | 1 |
| dmel | 1 |
| dmau, dsec, dsim | 1 |

## Notes

### Competing Interest Statement

The authors have declared no competing interest.

## References

[1] Adrian M Altenhoff et al. “OMA orthology in 2021: website overhaul, conserved isoforms, ancestral gene order and more”. In: Nucleic Acids Research 49.D1 (Nov. 2020), pp. D373–D379.

[2] Adrian M. Altenhoff et al. “OMA standalone: orthology inference among public and custom genomes and transcriptomes”. In: Genome Research 29.7 (June 2019), pp. 1152–1163.

[3] Adrian M. Altenhoff et al. “Resolving the Ortholog Conjecture: Orthologs Tend to Be Weakly, but Significantly, More Similar in Function than Paralogs”. In: PLoS Computational Biology 8.5 (May 2012). Ed. by Jonathan A. Eisen, e1002514.

[4] Stephen F. Altschul, Raymond J. Carroll, and David J. Lipman. “Weights for data related by a tree”. In: Journal of Molecular Biology 207.4 (June 1989), pp. 647–653.

[5] Christiam Camacho et al. “BLAST+: architecture and applications”. In: BMC Bioinformatics 10.1 (Dec. 2009).

[6] S. Capella-Gutierrez, J. M. Silla-Martinez, and T. Gabaldon. “trimAl: a tool for automated alignment trimming in large-scale phylogenetic analyses”. In: Bioinformatics 25.15 (June 2009), pp.1972–1973.

[7] J. Castresana. “Selection of Conserved Blocks from Multiple Alignments for Their Use in Phylogenetic Analysis”. In: Molecular Biology and Evolution 17.4 (Apr. 2000), pp. 540–552.

[8] F. Chen. “OrthoMCL-DB: querying a comprehensive multi-species collection of ortholog groups”. In: Nucleic Acids Research 34.90001 (Jan. 2006), pp. D363–D368.

[9] Drosophila 12 Genomes Consortium. “Evolution of genes and genomes on the Drosophila phylogeny”. In: Nature 450.7167 (Nov. 2007), pp. 203–218.

[10] Salvatore Cosentino and Wataru Iwasaki. “SonicParanoid: fast, accurate and easy orthology inference”. In: Bioinformatics 35.1 (July 2018). Ed. by Russell Schwartz, pp. 149–151.

[11] Th Dobzhansky. “Drosophila miranda, a new species”. In: Genetics 20.4 (July 1935), pp. 377–391.

[12] Michael P. Dunne and Steven Kelly. “OMGene: mutual improvement of gene models through optimisation of evolutionary conservation”. In: BMC Genomics 19.1 (Apr. 2018).

[13] Michael P. Dunne and Steven Kelly. “OrthoFiller: utilising data from multiple species to improve the completeness of genome annotations”. In: BMC Genomics 18.1 (May 2017).

[14] R. C. Edgar. “MUSCLE: multiple sequence alignment with high accuracy and high throughput”. In: Nucleic Acids Research 32.5 (Mar. 2004), pp. 1792–1797.

[15] David M. Emms and Steven Kelly. “OrthoFinder: phylogenetic orthology inference for comparative genomics”. In: Genome Biology 20.1 (Nov. 2019).

[16] David M. Emms and Steven Kelly. “OrthoFinder: solving fundamental biases in whole genome comparisons dramatically improves orthogroup inference accuracy”. In: Genome Biology 16.1 (Aug. 2015).

[17] A. J. Enright. “An efficient algorithm for large-scale detection of protein families”. In: Nucleic Acids Research 30.7 (Apr. 2002), pp. 1575–1584.

[18] J. Felsenstein and G. A. Churchill. “A Hidden Markov Model approach to variation among sites in rate of evolution”. In: Molecular Biology and Evolution 13.1 (Jan. 1996), pp. 93–104.

[19] Shaohong Feng et al. “Dense sampling of bird diversity increases power of comparative genomics”. en. In: Nature 587.7833 (Nov. 2020), pp. 252–257.

[20] Walter M. Fitch. “Distinguishing homologous from analogous proteins”. In: Systematic Zoology 31.2 (June 1970), pp. 99–113.

[21] Robert D. Fleischmann et al. “Whole-Genome Random Sequencing and Assembly of *Haemophilus influenzae* Rd”. In: Science 269.5223 (July 1995), pp. 496–512.

[22] Adam Frankish et al. “Comparison of GENCODE and RefSeq gene annotation and the impact of reference geneset on variant effect prediction”. In: BMC Genomics 16.S8 (June 2015).

[23] Michael Y Galperin et al. “COG database update: focus on microbial diversity, model organisms, and widespread pathogens”. In: Nucleic Acids Research 49.D1 (Nov. 2020), pp. D274–D281.

[24] A. Goffeau et al. “Life with 6000 Genes”. In: Science 274.5287 (Oct. 1996), pp. 546–567.

[25] L Sian Gramates et al. “FlyBase: a guided tour of highlighted features”. In: Genetics 220.4 (Mar. 2022). Ed. by V Wood.

[26] Charles R. Harris et al. “Array programming with NumPy”. In: Nature 585.7825 (Sept. 2020), pp. 357–362.

[27] Javier Herrero et al. “Ensembl comparative genomics resources”. In: Database 2016 (2016), bav096.

[28] Jaime Huerta-Cepas et al. “eggNOG 5.0: a hierarchical, functionally and phylogenetically annotated orthology resource based on 5090 organisms and 2502 viruses”. In: Nucleic Acids Research 47.D1 (Nov. 2018), pp. D309–D314.

[29] John D. Hunter. “Matplotlib: A 2D Graphics Environment”. In: Computing in Science &amp: Engineering 9.3 (2007), pp. 90–95.

[30] L. J. Jensen et al. “eggNOG: automated construction and annotation of orthologous groups of genes”. In: Nucleic Acids Research 36.Database (Dec. 2007), pp. D250–D254.

[31] Mateusz Kaduk and Erik Sonnhammer. “Improved orthology inference with Hieranoid 2”. In: Bioinformatics (Jan. 2017), btw774.

[32] K. Katoh and D. M. Standley. “MAFFT Multiple Sequence Alignment Software Version 7: Improvements in Performance and Usability”. In: Molecular Biology and Evolution 30.4 (Jan. 2013), pp. 772–780.

[33] Kazutaka Katoh and Daron M. Standley. “A simple method to control over-alignment in the MAFFT multiple sequence alignment program”. In: Bioinformatics 32.13 (Feb. 2016), pp. 1933–1942.

[34] Bernard Y Kim et al. “Highly contiguous assemblies of 101 drosophilid genomes”. In: eLife 10 (July 2021).

[35] Anders Krogh and Søren Kamaric Riis. “Hidden Neural Networks”. In: Neural Computation 11.2 (Feb. 1999), pp. 541–563.

[36] J.-L. Da Lage et al. “A phylogeny of Drosophilidae using the Amyrel gene: questioning the Drosophila melanogaster species group boundaries”. In: Journal of Zoological Systematics and Evolutionary Research 45.1 (Feb. 2007), pp. 47–63.

[37] S. Q. Le and O. Gascuel. “An Improved General Amino Acid Replacement Matrix”. In: Molecular Biology and Evolution 25.7 (Apr. 2008), pp. 1307–1320.

[38] Li Li, Christian J. Stoeckert, and David S. Roos. “OrthoMCL: Identification of Ortholog Groups for Eukaryotic Genomes”. In: Genome Research 13.9 (Sept. 2003), pp. 2178–2189.

[39] Benjamin Linard et al. “OrthoInspector 2.0: Software and database updates”. In: Bioinformatics 31.3 (Oct. 2014), pp. 447–448.

[40] Benjamin Linard et al. “OrthoInspector: comprehensive orthology analysis and visual exploration”. In: BMC Bioinformatics 12.1 (Jan. 2011).

[41] Martín Abadi et al. TensorFlow: Large-Scale Machine Learning on Heterogeneous Systems. Software available from tensorflow.org. 2015.

[42] Wes McKinney. “Data Structures for Statistical Computing in Python”. In: Proceedings of the Python in Science Conference. SciPy, 2010.

[43] Danny E Miller et al. “Highly Contiguous Genome Assemblies of 15 *Drosophila* Species Generated Using Nanopore Sequencing”. In: G3 Genes|Genomes|Genetics 8.10 (Oct. 2018), pp. 3131–3141.

[44] Yannis Nevers et al. “OrthoInspector 3.0: open portal for comparative genomics”. In: Nucleic Acids Research 47.D1 (Oct. 2018), pp. D411–D418.

[45] Yannis Nevers et al. “The Quest for Orthologs orthology benchmark service in 2022”. In: Nucleic Acids Research 50.W1 (May 2022), W623–W632.

[46] Lam-Tung Nguyen et al. “IQ-TREE: A Fast and Effective Stochastic Algorithm for Estimating Maximum Likelihood Phylogenies”. In: Molecular Biology and Evolution 32.1 (Nov. 2014), pp. 268–274.

[47] Alex Nord et al. “Splice-Aware Multiple Sequence Alignment of Protein Isoforms”. In: Proceedings of the 2018 ACM International Conference on Bioinformatics, Computational Biology, and Health Informatics. ACM, Aug. 2018.

[48] Cédric Notredame, Desmond G Higgins, and Jaap Heringa. “T-coffee: a novel method for fast and accurate multiple sequence alignment 1 1Edited by J. Thornton”. In: Journal of Molecular Biology 302.1 (Sept. 2000), pp. 205–217.

[49] Martin A. Nowak et al. “Evolution of genetic redundancy”. In: Nature 388.6638 (July 1997), pp. 167–171.

[50] Darren J. Obbard et al. “Estimating Divergence Dates and Substitution Rates in the Drosophila Phylogeny”. In: Molecular Biology and Evolution 29.11 (Aug. 2012), pp. 3459–3473.

[51] Susumu Ohno. Evolution by Gene Duplication. Springer Berlin Heidelberg, 1970.

[52] Cinta Pegueroles, Steve Laurie, and M. Mar Albà. “Accelerated Evolution after Gene Duplication: A Time-Dependent Process Affecting Just One Copy”. In: Molecular Biology and Evolution 30.8 (Apr. 2013), pp. 1830–1842.

[53] Maido Remm, Christian E.V. Storm, and Erik L.L. Sonnhammer. “Automatic clustering of orthologs and in-paralogs from pairwise species comparisons”. In: Journal of Molecular Biology 314.5 (Dec. 2001), pp. 1041–1052.

[54] Arang Rhie et al. “Towards complete and error-free genome assemblies of all vertebrate species”. In: Nature 592.7856 (Apr. 2021), pp. 737–746.

[55] The textitC. elegans Sequencing Consortium. “Genome Sequence of the Nematode *C. elegans*: A Platform for Investigating Biology”. In: Science 282.5396 (Dec. 1998), pp. 2012–2018.

[56] Fabian Sievers and Desmond G. Higgins. “Clustal Omega for making accurate alignments of many protein sequences”. In: Protein Science 27.1 (Oct. 2017), pp. 135–145.

[57] Patricia S. Soria, Kriston L. McGary, and Antonis Rokas. “Functional Divergence for Every Paralog”. In: Molecular Biology and Evolution 31.4 (Jan. 2014), pp. 984–992.

[58] Roman L. Tatusov, Eugene V. Koonin, and David J. Lipman. “A Genomic Perspective on Protein Families”. In: Science 278.5338 (Oct. 1997), pp. 631–637.

[59] Françoise Thibaud-Nissen et al. “The NCBI Handbook”. In: National Center for Biotechnology Information, 2013. Chap. Eukaryotic Genome Annotation Pipeline.

[60] Gregg W. C. Thomas et al. “Gene content evolution in the arthropods”. In: Genome Biology 21.1 (Jan. 2020).

[61] Clément-Marie Train et al. “Orthologous Matrix (OMA) algorithm 2.0: more robust to asymmetric evolutionary rates and more scalable hierarchical orthologous group inference”. In: Bioinformatics 33.14 (July 2017), pp. i75–i82.

[62] Pauli Virtanen et al. “SciPy 1.0: fundamental algorithms for scientific computing in Python”. In: Nature Methods 17.3 (Feb. 2020), pp. 261–272.

[63] Brian M. Wiegmann et al. “Episodic radiations in the fly tree of life”. In: Proceedings of the National Academy of Sciences 108.14 (Mar. 2011), pp. 5690–5695.

[64] Ziheng Yang. “Maximum likelihood phylogenetic estimation from DNA sequences with variable rates over sites: Approximate methods”. In: Journal of Molecular Evolution 39.3 (Sept. 1994), pp. 306–314.

[65] Evgeny M Zdobnov et al. “OrthoDB in 2020: evolutionary and functional annotations of orthologs”. In: Nucleic Acids Research 49.D1 (Nov. 2020), pp. D389–D393.

