## Supplementary material for "Leveraging genomic redundancy to improve inference and alignment of orthologous proteins": README.txt

The following folders are included in the supplemental files. The folders are listed in order of their creation by the orthology inference pipeline.

clusters/: Orthologous groups generated by 4-clique percolation with paralogs added.
align_fastas/: The aligned sequences for each cluster in clusters4.tsv.
get_repseqs/: The aligned representative sequences for each alignment in align_fastas/.
insertion_trim/: The alignments of single copy orthologs in get_repseqs/ with the insertions identified by the insertion phylo-HMM removed.
missing_trim/: Missing segments identified by the missing phylo-HMM for each sequence in each single copy orthologous group. Missing segments are given as Python slices in the following form: start0-stop0,start1-stop1,...,startn-stopn. If the slices field is empty, then no missing segments were identified in that sequence.
